# Fibronectin Composition and Transglutaminase 2 Cross-linking Cooperatively Regulate Ovarian Cancer Cell Adhesion in ECM-Mimetic Constructs

**DOI:** 10.1101/2025.05.13.653765

**Authors:** Ning Yang, Ali Abbaspour, James M. Considine, Stephanie M. McGregor, Erin G. Brooks, Alexandra Naba, Kristyn S. Masters, Pamela K. Kreeger

## Abstract

The extracellular matrix (ECM) plays a crucial role in tumor progression. Here, we analyzed collagen I and cellular fibronectin (cFN) in normal omentum and metastatic omentum from high-grade serous ovarian cancer (HGSOC). The levels of both proteins were significantly elevated and collagen I fibers were significantly thicker in HGSOC metastases. Moreover, the ECM cross-linking enzyme transglutaminase 2 (TG2) was increased in omental metastases, where it is enzymatically active in the extracellular environment. This information was used to develop ECM constructs recapitulating these key changes, alone and in combination, to investigate their impact on HGSOC cell adhesion. To our knowledge, this is the first report using TG2 as a cross-linking agent to generate constructs from multiple ECM components. Low levels of HGSOC cell adhesion were observed on colIagen-only (coll) gels, while inclusion of cFN or plasma fibronection (pFN) increased cell adhesion. TG2-mediated cross-linking of colI/cFN hydrogels promoted HGSOC cell adhesion, while cross-linking of coll/pFN had no effect. Cell adhesion was dependent on ligand identity and fiber diameter. When fiber thickness was held constant, the inclusion of cFN led to greater HGSOC cell adhesion relative to pFN or coll, due to interactions of β1 integrins with the EDA and RGD domains of cFN. Meanwhile, when gel composition was held constant, HGSOC cell adhesion increased as fiber thickness was increased through modifications to gelation temperature. Combined, our results demonstrate how ECM changes associated with omental metastasis can support tumor progression and provide insights into methods to tailor biomaterials to support cell adhesion.

## Introduction

The critical role of the extracellular matrix (ECM) in many developmental and pathological processes has led to the development of a wide range of biomaterials to simulate its complex structure and biochemical cues [1, 2]. Many of these ECM-based biomaterials utilize fibrillar collagen (primarily collagen I), a key structural component of the ECM that is commonly overexpressed and undergoes architectural changes in diseases such as fibrosis and cancer. Both the presence of collagen fibers and their arrangement can influence disease progression. For example, in an *in vitro* model of breast cancer, tumor cells were unable to invade the surrounding matrix in the absence of collagen fibers [3]. The alignment of collagen I fibers is also a key microarchitectural signature of breast cancer and is linked with increased cancer invasion and malignancy [4].

*In vivo*, many regulatory processes impact the physical and mechanical characteristics of collagen fibers, including post-translational modifications (*e.g.*, cross-linking, proteolytic cleavage) and protein-protein interactions with other ECM proteins or with cell-surface receptors [5]. *In vitro*, changes in collagen fiber architecture can be mimicked through various methods, including increasing fiber thickness by adjusting the gelation temperature [6] and aligning fibers using magnetic beads, microfluidic channels, mechanical stretching, or collagen cross-linking enzymes [7–12]. To date, these efforts have not included other components of the ECM that interact with collagen fibers *in vivo*, such as fibronectin (FN). FN and collagen I colocalize in many tissues. Their interaction influences the assembly and organization of ECM fibers, altering both the biological and physical properties of the ECM and impacting cell behavior [13, 14].

Collagen binds to the gelatin-binding domain of FN, which facilitates the assembly of collagen fibrils [15]. Moreover, enzymes such as transglutaminase 2 (TG2) can regulate the interaction between collagen and FN. TG2 exhibits Ca^2+^-dependent transamidase activity, resulting in covalent cross-links between substrate proteins through the creation of isopeptide bonds between the γ-carboxamide group of a glutamine residue and the ε-amino group of a lysine residue [16]. TG2 is implicated in cross-linking of adjacent ECM protein polymers including FN and collagen *in vivo* [17]. In addition, TG2 has been used as a nontoxic biological cross-linker of self-assembled colI hydrogel to create cross-linked gels with increased thermal stability and altered fibril structure [12].

The present study examined whether TG2 could be used to create fibrillar ECM scaffolds that better mimic the tumor stroma by incorporating collagen and FN. We focused on metastatic ovarian cancer, a particularly aggressive type of cancer that represents the 5^th^ leading cause of cancer death in women in the United States. Previous studies have shown that both collagen [18] and FN [19] levels are drastically elevated in the metastatic ovarian cancer microenvironment and particularly in the omentum, the most common site of ovarian cancer metastasis. Here, using human omental samples from female decedents without cancer and omental metastases of patients with high-grade serous ovarian cancer (HGSOC), we first examined the changes in fibrillar collagen, cFN, and TG2 levels and found that TG2 was enzymatically active in the extracellular environment. We then used this information to develop biomaterials that mimic the changes observed in metastatic tissues and investigated their impact on ovarian cancer cell adhesion.

## Materials and Methods

All reagents were purchased from Thermo Fisher Scientific unless noted otherwise.

### Cell culture

Of the three HGSOC cell lines used in the present study, OVCAR3 and OVCAR4 were purchased from American Type Culture Collection (ATCC) while OVCAR8 was obtained from NCI 60 panel (NIH). The cell lines were authenticated by the Translational Research Initiatives in Pathology (TRIP) Laboratory at the University of Wisconsin-Madison using human short tandem repeat analysis. Cells were maintained in 1:1 Medium 199 (with Earle’s salts and L-glutamine, Sigma-Aldrich) and MCDB 105 medium (Sigma-Aldrich) containing 15% heat-inactivated fetal bovine serum (HI-FBS) and 1% penicillin-streptomycin. Cell experiments were conducted in serum-free medium (SFM).

### Picrosirius red (PSR) staining of collagen

De-identified, formalin-fixed, paraffin-embedded (FFPE) omental samples from patients with stage III/IV HGSOC were obtained from archived pathology samples through a protocol approved by the Institutional Review Board (IRB #2016-1176) at the University of Wisconsin-Madison (Supplemental Table 1). Normal omental samples were collected from deceased females with no history of tumor during autopsy and were classified as not human subjects research by the IRB. To examine collagen levels in normal and HGSOC omental tissues, 5 μm sections were prepared by the TRIP lab, deparaffinized with SafeClear II xylene substitute, rehydrated in gradients of ethanol, and then incubated with PSR solution at RT for 1 h. The PSR solution was prepared by dissolving 0.5 g of Direct Red 80 (Sigma-Aldrich) in 500 mL of saturated picric acid (Sigma-Aldrich). Subsequently, sections were washed with 0.5% (v/v) acetic acid in water, dehydrated in 100% ethanol, cleared in xylene, and mounted with Richard-Allan toluene-based mounting medium. To determine fibrillar collagen levels in the omental samples, the stained sections were imaged on a Zeiss Axio Observer.Z1 inverted microscope with a Plan-Apochromat 20x/0.8 Ph2 M27 objective using the filter set for mCherry (Zeiss), and the mean fluorescence intensity (MFI) was measured using ImageJ (NIH). To determine the width of collagen fibers in the omental samples, the stained sections were imaged using a Nikon AXR laser scanning confocal microscope with a PLAN APO Lambda D 60X/0.13 oil objective and a TRITC filter set (570-620 nm, emission, Nikon) and analyzed using the CT-FIRE and CURVE-Align software package (loci.wisc.edu/software/ctFIRE, v.2.0b & loci.wisc.edu/software/curvealign, v.5.0) [20]. For each omental sample, the mean value of at least five analyzed images was reported.

### Immunohistochemical staining of fibronectin and TG2

Levels of cellular fibronectin (cFN), TG2 and TG2-mediated cross-linking in normal and HGSOC omental tissues were examined by immunohistochemical staining. Omental sections were deparaffinized, rehydrated in graded ethanol series, and subjected to antigen retrieval in antigen unmasking solution (Vector Laboratories) at 80°C for 1 h. The sections were then blocked with 5% goat serum in PBS and incubated with primary antibodies diluted in blocking buffer at 4°C overnight. The primary antibodies used in the present study included EDA+ FN (ab6328; Abcam, Waltham, MA; 10 μg/mL), EDB+ FN (MBS488254, MyBioSource, 10 μg/mL), TG2 (ab421, Abcam, 5 μg/mL), and N-epsilon gamma-glutamyl lysine (ab424, Abcam, 10 μg/mL). Secondary antibodies (Alexa Fluor 647-conjugated goat anti-rabbit IgG, A-21245, 10 μg/mL; Alexa Fluor 647-conjugated goat anti-mouse IgG, A-21236, 10 μg/mL; and Cy5-conjugated Affinipure goat anti-mouse IgM, 115-175-075, Jackson Immuno Research Labs, 7.5 μg/mL) were applied at room temperature (RT) for 1 h. Slides were mounted using ProLong Diamond Antifade with DAPI.

In all staining, a control that omitted the primary antibody was included to determine the nonspecific background fluorescence. Images were obtained using a Zeiss Axio Observer.Z1 inverted microscope with an AxioCam 506 mono camera and a Plan-Apochromat 20x/0.8 Ph2 M27 or a LD Plan-Neofluar 40x/0.6 Korr M27 objective. Protein levels were evaluated using ImageJ by measuring MFI with background fluorescence subtracted.

### Preparation of ECM-mimetic constructs

All ECM constructs except for the ones used for dynamic mechanical analysis (DMA) were prepared in 96-well 3D µ-Plates (89646, ibidi). Briefly, FibriCol collagen (5133, Advanced Biomatrix,) was neutralized with 0.1 N NaOH/10X PBS and mixed with a solution of human fibronectin, either from human foreskin fibroblasts (cFN; F2518, Sigma-Aldrich) or plasma (pFN; 5080, Advanced Biomatrix), or SFM to achieve various final concentrations of 2.5 mg/mL for collagen and 0 or 60 μg/mL for fibronectin. Ten microliters of the mixture were added to each well of the plate and incubated at 37°C for 1 h to allow for polymerization. Enzymatic cross-linking was achieved by immersing the gels in 45 μL of SFM containing 1 mM DTT, 2.5 mM CaCl_2_, 25 mM MOPS, and 1.2 units/mL TG2 (23595, Cayman Chemical) and incubating at 37°C overnight. The enzymatic activity of each lot of TG2 was determined by a transglutaminase activity assay (CS1070, Sigma*-*Aldrich). The gels were washed 3 times with SFM before seeding cells to remove TG2 from the ECM constructs.

To alter fiber width, the colI and colI/cFN constructs were also prepared under slow gelation conditions at lower temperature. Specifically, the colI and the colI/cFN mixtures were added to a µ-Plate that was precooled at 4°C. The mixtures were then incubated at 4°C overnight, followed by incubation at RT for 1.5 h and 37°C for 1 h (referred to as 4C O/N). The colI/cFN construct was also prepared by incubation at 4°C for 1.5 h, followed by RT for 1.5 h and 37°C for 1 h (referred to as 4C 1.5 h).

### Measurement of fiber architecture using collagen-binding adhesion protein 35 (CNA35-EGFP)

To analyze collagen fibers in the ECM constructs, gels were blocked with 5% bovine serum albumin (BSA) in PBS at 4°C overnight and then incubated with 50 μM enhanced CNA35-EGFP at 37°C overnight. The CNA35-EGFP, a fluorescently tagged collagen-binding protein that preferentially labels fibrillar collagen, was expressed and purified from *E. coli* as previously described [21]. The plasmid encoding CNA35-EGFP, pET28a-EGFP-CNA35, was obtained from Maarten Merkx (Addgene plasmid #61603). To minimize the background fluorescence caused by unbound CNA35-EGFP trapped in the gel, the gels were washed with 0.1% BSA in PBS for at least 3 days (washing buffer was changed every 2-3 hours) before being imaged with a Nikon AXR laser scanning confocal microscope at 100X magnification using the FITC (500-550 nm,

emission) filter set. For each individual gel, at least three Z-stack images were taken at 5-μm intervals through the bottom 50 µm of the gels. The top, middle and bottom images of each Z-stack were analyzed using CT-FIRE and the width of the Col I fibers was calculated as the mean of the three analyzed images per sample.

### In vitro HGSOC cell adhesion assay

HGSOC cells were seeded onto different ECM constructs in a 96-well µ*-*Plate at a density of 2000 cells/well in SFM and allowed to adhere for 30 min. Unattached cells were removed by washing the wells twice with PBS. Cells that had adhered to the constructs were fixed with 4% paraformaldehyde (PFA; Electron Microscopy Sciences) for 30 min, stained with Hoechst 33342 at 1:500, and imaged on a Zeiss Axio Observer.Z1 inverted microscope with a Plan-Apochromat 5x/0.16 air objective and DAPI filter. At least 3 wells were seeded for each condition. One image was captured per well at low (5X) magnification in the center of the well. The images covered an area of 0.05 cm^2^, accounting for 40% of the growth area of the wells. The percentage of adherent cells was calculated by converting the number of adherent cells per image to total number of adherent cells per well and then dividing by the number of cells initially added to each well.

### Analysis of ascites samples for TG2

Ascites samples were collected from female patients with benign conditions or stage III/IV HGSOC during surgery. The University of Wisconsin Carbone Cancer Center Translational Science BioCore acted as an honest broker under IRB #2016–0934, obtaining informed consent from all participants and de-identifying samples before providing to the research team. Cells were removed from the ascites by centrifuging at 300 x g for 10 min. The amount and enzymatic activity of TG2 in the fluid portion of the ascites were determined using the Human TGM2 ELISA Kit (MBS2507350, MyBioSource) and the Transglutaminase Assay Kit (CS1070, Sigma*-*Aldrich), respectively, in accordance with the manufacturers’ instructions.

### Compartmental protein extraction and western blot analysis for TG2

Flash frozen human HGSOC omental metastases were procured from debulking surgeries. The University of Wisconsin Carbone Cancer Center Translational Science BioCore acted as an honest broker under IRB #2016–0934, obtaining informed consent from all participants and de-identifying samples before providing to the research team. In brief, samples were homogenized using a Bead Ruptor (Omni International) at 4°C. Enrichment of insoluble proteins was achieved through sequential extraction using the Subcellular Protein Fractionation Kit for Tissue (Thermo Scientific) and a protocol previously published (https://dx.doi.org/10.17504/protocols.io.kxygx94b4g8j/v1). Proteins were monitored by western blot analysis using a serum containing anti-fibronectin antibodies (rabbit 297-1, used at a 1:5000 dilution) kindly gifted by Dr. Richard Hynes (MIT) and a rabbit anti-transglutaminase 2 antibody (ab421, Abcam, 2.5 μg/mL).

### Immunofluorescence staining for TG2-mediated cross-linking in the ECM constructs

To determine whether TG2 treatment induced protein cross-linking in the ECM constructs, the colI/cFN gels were blocked with 5% BSA in PBS at 4°C overnight, incubated with anti-N-epsilon gamma-glutamyl lysine antibody (ab424, Abcam, 10 μg/mL) in the blocking buffer at 4°C overnight, and then incubated with Cy5-conjugated Affinipure goat anti-mouse IgM (115-175-075, Jackson Immuno Research Labs, 7.5 μg/mL) at RT for 1 h. To minimize background fluorescence caused by antibody trapped in the gel, gels were washed with 0.1% BSA in PBS for at least 3 days (washing buffer was changed every 2-3 h) before being imaged on the Zeiss Axio Observer.Z1 inverted microscope with a Plan-Apochromat 20x/0.8 Ph2 M27 objective and the Cy5 filter. The level of TG2-mediated protein cross-linking was evaluated using ImageJ by measuring MFI after subtracting the background fluorescence in no-primary antibody controls.

### DMA analysis

The stiffness of the ECM constructs was examined by DMA analysis. The ECM constructs were prepared as discs in 8 mm x 2 mm silicone molds. Mechanical tests were performed using an RSA III dynamic mechanical analyzer (TA Instruments). Prior to starting any tests, an initial minimum force preloading condition of ∼1 g was applied to the gels. Dynamic strain sweep tests were performed at room temperature at constant frequency (1 Hz) and oscillating strain ranging from 0.01% to 50%. Following the DMA evaluation, we identified the storage modulus (G’) and loss modulus (G’’) and the complex modulus (G) was obtained using the following formula: 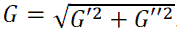.

### Immunocytochemistry for activated β1 integrin

HGSOC cells were fixed in 4% PFA for 30 min, permeabilized with 5% BSA/0.3% Triton X-100 in PBS at RT for 1 h, and blocked with 5% BSA in PBS at 4°C overnight. The cells were then incubated with primary antibody against activated β1 integrin (MAB2079Z, Sigma-Aldrich, 1:100) at 4°C overnight, followed by incubation with Alexa Fluor 647-conjugated goat anti-mouse IgG (1:200) at RT for 1 h. Cells were counter-stained with Hoechst 33342 (1:500) at RT for 20 min prior to imaging using a Zeiss Axio Observer.Z1 inverted microscope with a Plan-Apochromat 20x/0.8 Ph2 M27 objective and Alexa 647/DAPI filter sets. Total fluorescence per cell was evaluated using ImageJ. Higher-magnification images were also obtained using a Nikon AXR laser scanning confocal microscope at 40X magnification using the DAPI (424-475nm, emission) and Cy5 (655-850nm, emission) filter sets.

### Integrin inhibition

HGSOC cells were trypsinized and resuspended in SFM at 40,000 cells/mL. Irigenin (34676, Cayman Chemical) was added to the cell suspension at a final concentration of 50 μM or Volociximab (NBP2-52680; Novus Biologicals) was added at 15 μg/mL. A vehicle control and an isotype control were prepared by using an equivalent volume of DMSO or rabbit IgG isotype control antibody (Thermo Fisher, 31235), respectively. After incubation for 30 min, 50 μL of the cell suspension was seeded onto each ECM construct and cell attachment was assessed as described above 30 min after seeding.

### Statistical analysis

All data are presented as mean ± SD. Statistical tests were performed in GraphPad Prism and are detailed in figure legends. p<0.05 was considered statistically significant.

## Results

### Thicker collagen fibers are observed in human HGSOC omental metastases and promote HGSOC cell adhesion

To design fibrillar ECM scaffolds that mimic the tumor stroma environment, we first characterized the ECM composition in HGSOC omental metastases compared to healthy omentum. Based on prior reports [19], the most abundant ECM protein in the omentum is collagen I. Therefore, we stained tissues (Supplemental Table 1) with PSR and found that while collagen I in the healthy omentum was organized as a lacy structure around the adipocytes, in metastatic tissue there were regions of very dense collagen with almost no adipocytes remaining (Fig. 1A). Quantification of PSR staining of these tissues revealed a significant increase in collagen in the metastatic omentum compared with healthy tissues (Fig. 1B).

**Fig. 1.**
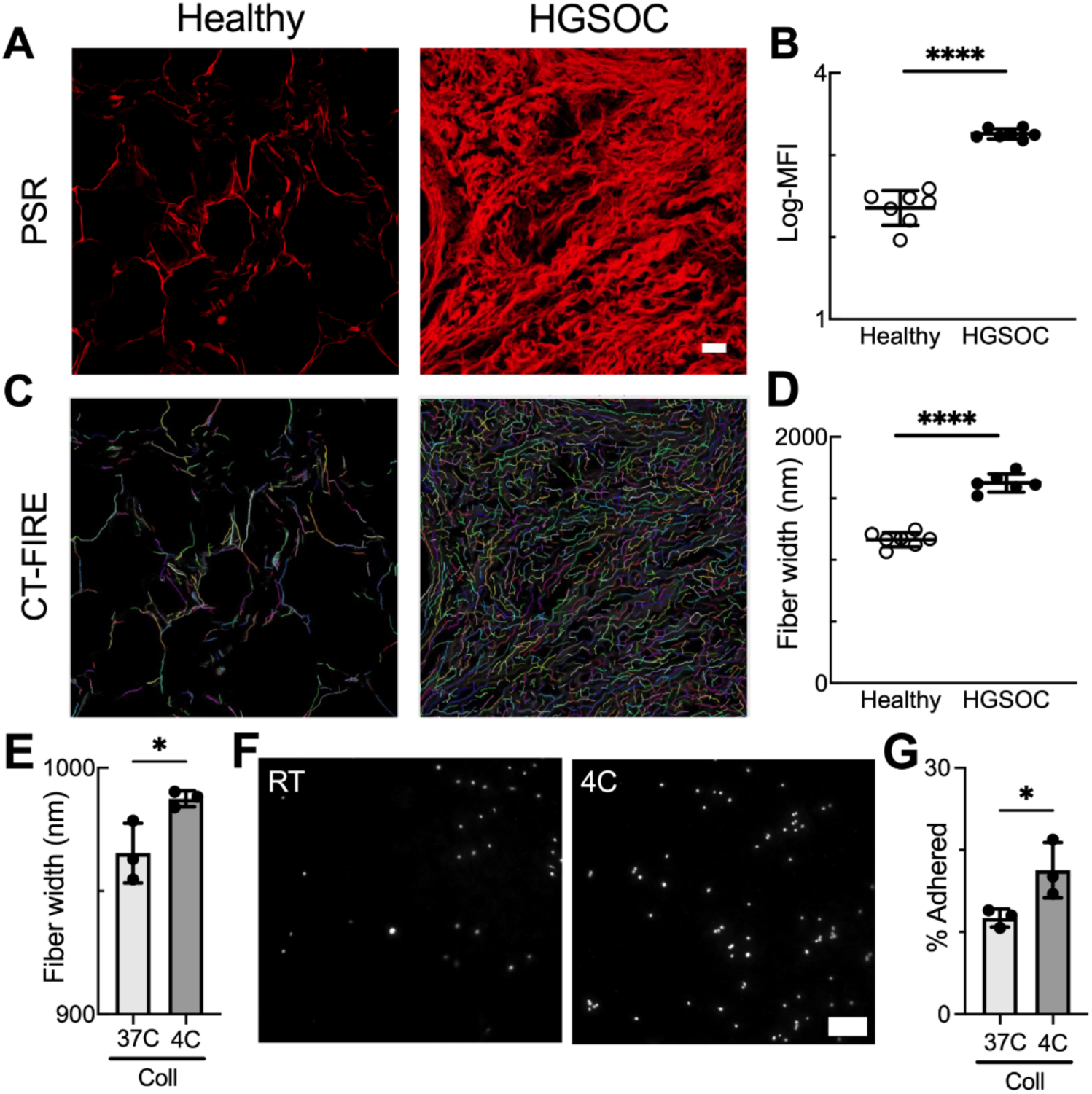
Collagen fibers are significantly thicker in human HGSOC omental metastases and thicker fibers support increased tumor cell attachment. **A,** Representative images of PSR staining of human omental samples. Scale bar = 20 μm. **B,** Quantification of mean fluorescent intensity (MFI) for samples in **(A)**. Each dot represents the average across at least 5 fields of view for an individual patient. **** indicates *p*<0.0001 by unpaired t-test with Welch’s correction. **C,** Representative CT-FIRE identification of PSR-stained fibers in human normal and metastatic omental samples. **D,** Mean fiber width of CT-FIRE identified fibers for samples in **(C)**. Each dot represents the average across at least 5 fields of view for an individual patient. **** indicates *p*<0.0001 by unpaired t-test. **E,** Mean fiber width of CT-FIRE identified fibers in coll gels formed at 37°C for one hour (37C) or 4°C overnight (4C). Each dot represents the average across at least 9 fields of view of an individual gel, * indicates p<0.05 by unpaired t-test. **F,** Representative images of Hoechst-stained OVCAR4 cells attached on coll gels. Scale bar = 200 μm. **G,** Quantification of OVCAR4 cells attached on coll gels. Each dot represents an individual gel, * indicates p<0.05 by unpaired t-test.

Moreover, analysis of confocal images of collagen fibers using CT-FIRE revealed that individual fiber width was significantly increased in the metastatic omentum compared with healthy omentum (Fig. 1C,D).

We have recently demonstrated that HGSOC adhesion is increased in the presence of fibrillar collagen relative to denatured gelatin for cells on biomaterial constructs [22]. Therefore, we next sought to examine the impact of collagen fiber thickening on HGSOC cell adhesion. We have previously determined that the concentration of collagen in grossly normal regions of a metastatic omentum is approximately 2.5 mg/mL [18]. We prepared constructs composed of 2.5 mg/mL coll and polymerized them for an hour at 37°C or overnight at 4°C followed by shorter incubations at room temperature and then 37°C. Fibers in these constructs were labeled with green fluorescent protein (EGFP)-tagged collagen-binding adhesion protein 35 (CNA35-EGFP), imaged by confocal microscopy and analyzed by CT-FIRE. Consistent with previous reports that temperature during gelation regulates collagen fiber width, gelation at 4°C overnight resulted in a significant increase in the mean fiber width (Fig. 1E, Fig. S1A). Three HGSOC cell lines (OVCAR3, OVCAR4 and OVCAR8) were seeded on the two constructs and assessed for adhesion 30 minutes after seeding. OVCAR3 and OVCAR4 had a significant increase in attachment on gels incubated at 4°C (Fig. 1F,G; Fig. S1B); OVCAR8 showed a similar trend but the difference was not statistically significant (Fig. S1B). These results suggest that thicker collagen fibers facilitate HGSOC cell adhesion.

### Fibronectin is increased in human HGSOC omental metastases but has a modest effect on HGSOC cell attachment

We next examined fibronectin in the omental samples, based on reports that the abundance of FN is significantly increased in omental tumors [19] and the known interactions between collagen and fibronectin in the ECM [23]. Cellular FN (cFN) is the form of FN most commonly found in the ECM and is produced by cells such as fibroblasts, endothelial cells and muscle cells [24]. Meanwhile, plasma FN (pFN) is secreted into the blood in a soluble and compact form by hepatocytes. Because cellular fibronectin (cFN) is strongly associated with the insoluble ECM, we focused on detection of cFN splice variants. We found that both the extra domain A and extra domain B splice variants of cFN (EDA-FN and EDB-FN [24]) were significantly elevated in metastatic omental tissues compared to healthy omental tissues (Fig. 2A-D).

**Fig. 2.**
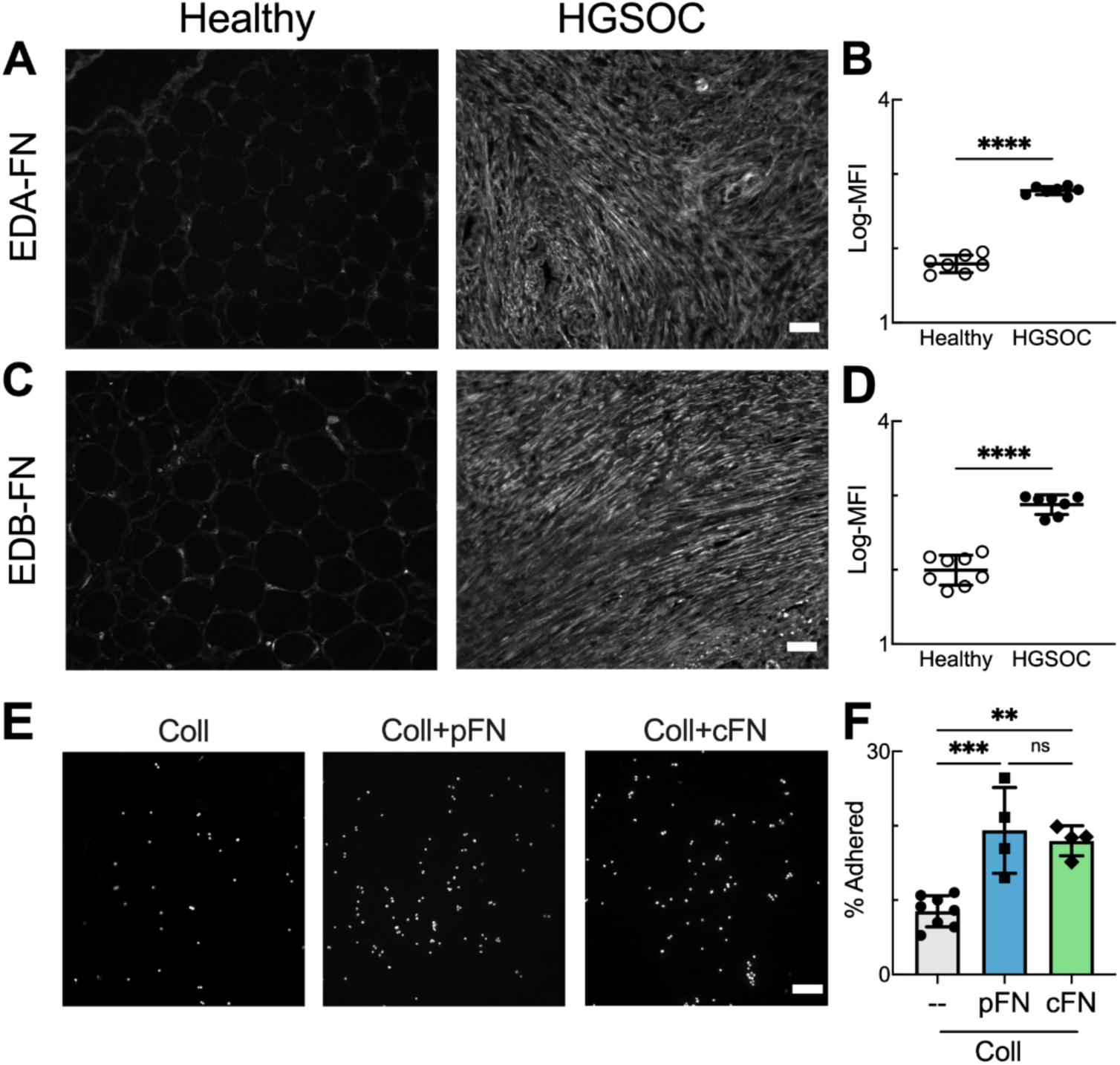
Fibronectin is increased in human HGSOC omental metastases and supports increased tumor cell attachment. **A,** Representative images of EDA-FN immunostained human omental samples. **B,** Quantification of MFI for samples in **(A)**. **C,** Representative images of EDB-FN immunostained human omental samples. **D,** Quantification of MFI for samples in **(C)**. Scale bar = 50 μm in **(A,C),** each dot represents the average across at least 5 fields of view for an individual patient in **(B,D)**. **** indicates *p*<0.0001 by unpaired t-test. **E,** Representative images of Hoechst-stained OVCAR4 cells attached on coll, coll+pFN, or coll+cFN. Scale bar = 200 μm. **F,** Quantification of OVCAR4 cells for conditions in **(E)**. Each dot represents an individual gel, ns indicates p>0.05, ** indicates p<0.01, *** indicates p<0.001 by Tukey.

Quantitative dot blot analysis of the omental samples showed that there was an approximately 6-fold increase in FN concentration in the metastatic omental tissues compared with the healthy omental tissues (∼60 μg/mL vs. 10 μg/mL) (Fig. S2A,B). Given the significant changes in cFN content in the metastatic omentum, we investigated how the inclusion of fibronectin would impact HGSOC cell adhesion. We first generated biomimetic constructs composed of 2.5 mg/mL coll and 60 μg/mL pFN (coll/pFN). While our tissue characterization focused on cFN, pFN contains the RGD integrin-binding domain and N-terminal collagen binding sites [24] and is widely used in biomaterials due to its affordability. Coll/pFN gels were allowed to polymerize at 37°C for 1 hour and cell adhesion examined at 30 minutes. While there was a significant increase in OVCAR4 adhesion on coll/pFN relative to coll alone (Fig. 2E,F), there was no effect on the adhesion of OVCAR3 or OVCAR8 cells (Fig. S2C). We then constructed gels of 2.5 mg/mL coll and 60 μg/mL cFN (coll/cFN) to examine if the inclusion of the two alternatively-spliced fibronectin type III domains EDA or EDB would result in different effects on cell adhesion. The *in vitro* adhesion assay showed a significant increase in OVCAR4 cell adhesion on colI/cFN relative to coll (Fig. 2E,F). A similar trend was observed with OVCAR3 cells, but no effect was seen with OVCAR8 (Fig. S2C). These results suggest that cFN increases HGSOC cell attachment when loosely incorporated into collagen gels, although the effect was cell-type specific.

### TG2 is enzymatically active extracellularly and induces protein cross-linking in the ECM of human HGSOC omental metastases

We next considered how ECM-modifying enzymes could alter the tumor microenvironment. The significant thickening of the collagen fibers observed in the metastatic omentum (Fig. 1) is consistent with the effects expected from enzymes such as TG2. TG2 is a multifunctional protein that exhibits conformation-dependent protein cross-linking activity and is located both in the ECM and intracellularly in the cytosol [16, 25, 26]. A variety of ECM proteins, including collagen and cFN, act as substrates for TG2; therefore, we investigated whether TG2 was present and active in the omental samples. Immunostaining revealed a markedly increased level of TG2 in metastatic omental tissues compared with the healthy omentum (Fig. 3A,B). Co-staining with cellular markers suggested that TG2 was expressed by both cancer cells and stromal cells in the metastatic omentum (Fig. S3). Since TG2 must be secreted and catalytically active in the extracellular environment to participate in ECM cross-linking, we next investigated if TG2 was present in the extracellular space in samples from patients with HGSOC or benign conditions (Supplemental Table 2). Benign conditions were used as healthy patients do not have large enough volumes of peritoneal fluid to allow for easy collection. TG2 was present in peritoneal ascites and was significantly elevated in stage III/IV HGSOC relative to patients with benign conditions (Fig. 3C). To determine if the extracellular TG2 was catalytically active, the ascites samples were analyzed using a transglutaminase assay kit, and activity was also found to be significantly elevated in stage III/IV HGSOC (Fig. 3D).

**Fig. 3.**
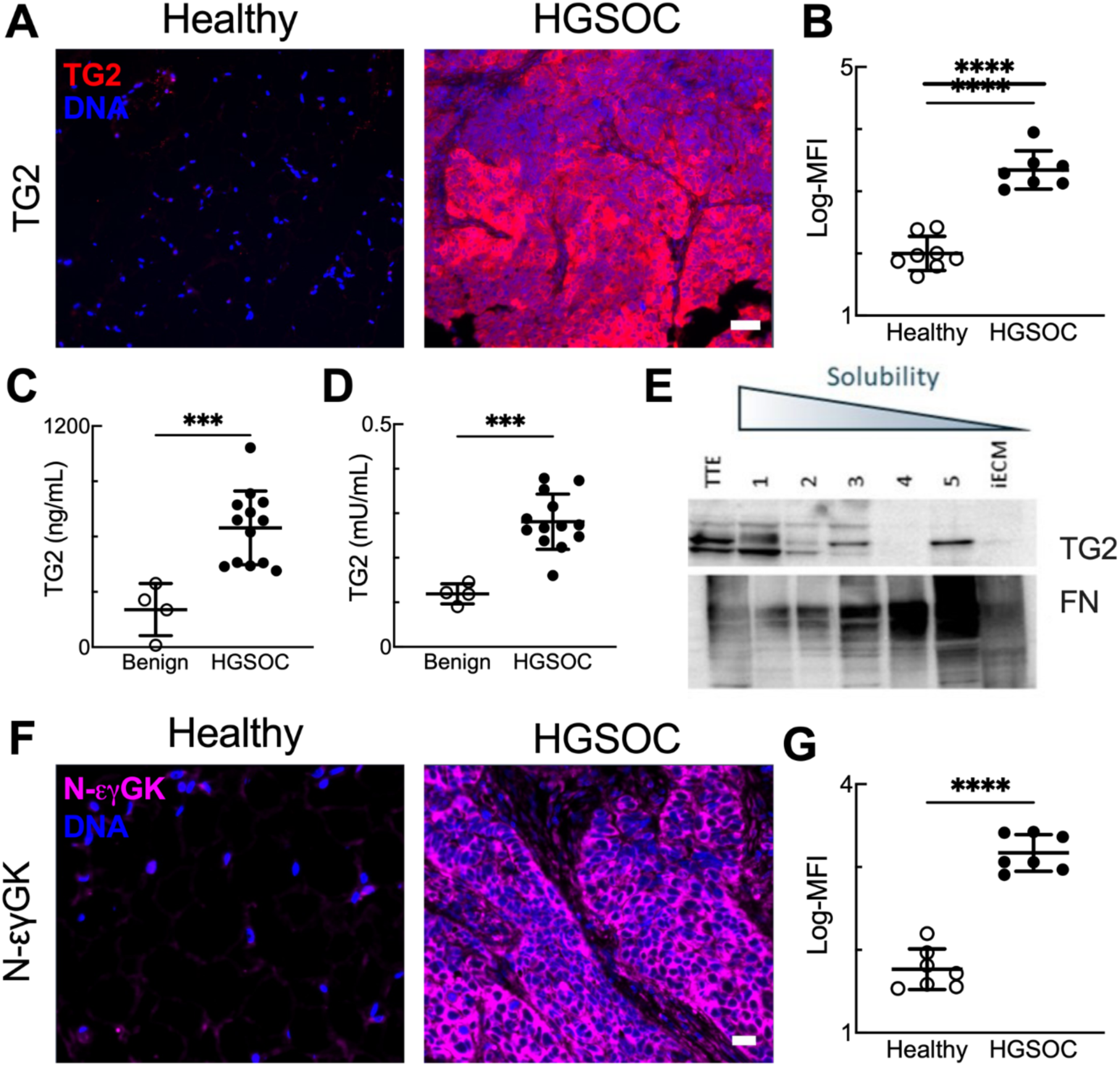
Transglutaminase-2 cross-linking is elevated in metastatic HGSOC. **A,** Representative images of TG2-immunostained human omental samples. Scale bar = 50 μm. **B,** Quantification of TG2 MFI for samples in **(A)**. Each dot represents the average across at least 5 fields of view for an individual patient. **** indicates *p*<0.0001 by unpaired t-test. **C, D** Comparison of TG2 level **(C)** and transglutaminase activity **(D)** between ascites samples collected from patients with benign conditions and patients with stage III/IV HGSOC. Each dot represents an individual patient, *** indicates p<0.001 by unpaired t-test. **E,** Western blot analysis of TG2 in different fractions of protein extracts from human HGSOC omental metastases. **F,** Representative images of N-εγGK-immunostained human omental samples. Scale bar = 20 μm. **G,** Quantification of N-εγGK MFI for samples in **(F)**. Each dot represents the average across at least 5 fields of view for an individual patient. **** indicates *p*<0.0001 by unpaired t-test.

To determine if TG2 was present within the omental ECM, metastatic omental tissue was subjected to subcellular fractionation and different fractions were assessed for TG2 by western blot (Supplemental Table 3). As expected, TG2 was found in the most soluble fractions, which are enriched for cytosolic proteins. However, a substantial amount of TG2 was also observed in the protein fractions that are highly insoluble (fraction 5 and the most insoluble ECM fraction), which are enriched for other ECM proteins such as FN (Fig. 3E; similar results were observed with a second sample).

When it adopts an open/extended conformation, TG2 catalyzes the cross-linking of proteins through the generation of N-epsilon-gamma glutamyl lysine isopeptide bonds (N-εγGK). Omental samples were immunostained for this cross-link (Fig. 3F,G) and the levels in metastatic omental tissues were found to be significantly elevated, confirming the enzymatic activity of TG2. Combined, these results indicate that TG2 is present extracellularly and is enzymatically active in HGSOC, resulting in elevated ECM cross-linking in the metastatic omentum.

### TG2-mediated ECM cross-linking promotes the attachment of HGSOC cells

We next sought to examine the impact of our observation of TG2-induced cross-linking on HGSOC cell adhesion in the context of collagen and fibronectin. We used the same formulations for coll, coll/pFN, and coll/cFN as in Fig. 2, with the addition of an incubation with 1.2 U/mL TG2 (Fig. 4A). Successful TG2-mediated cross-linking of the *in vitro* constructs was confirmed by a significant increase in N-εγGK immunostaining in the TG2-treated constructs (Fig. 4B,C). Cross-linked fibers are expected to be stiffer [27]; consistent with the observed N-εγGK cross-links, TG2-treated gels were nearly 2.5-fold stiffer than their uncross-linked counterparts (Fig. 4D). All constructs treated with TG2 had virtually the same stiffness, suggesting a similar level of protein cross-linking regardless of the presence of FN (Fig. S4A). HGSOC cells were seeded on these constructs and assessed for adhesion at 30 min after seeding (Fig. 4E; Fig. S4B). OVCAR4 and OVCAR8 had significantly increased attachment on coll+TG2 or coll/pFN+TG2 relative to coll or coll/pFN, respectively. More strikingly, all three cell lines showed significantly higher attachment on the colI/cFN+TG2 construct compared to all direct comparisons (coll+TG2, coll/pFN+TG2, coll/cFN). In summary, TG2-mediated cross-linking of the ECM promoted HGSOC cell adhesion and thus might support HGSOC progression.

**Fig. 4.**
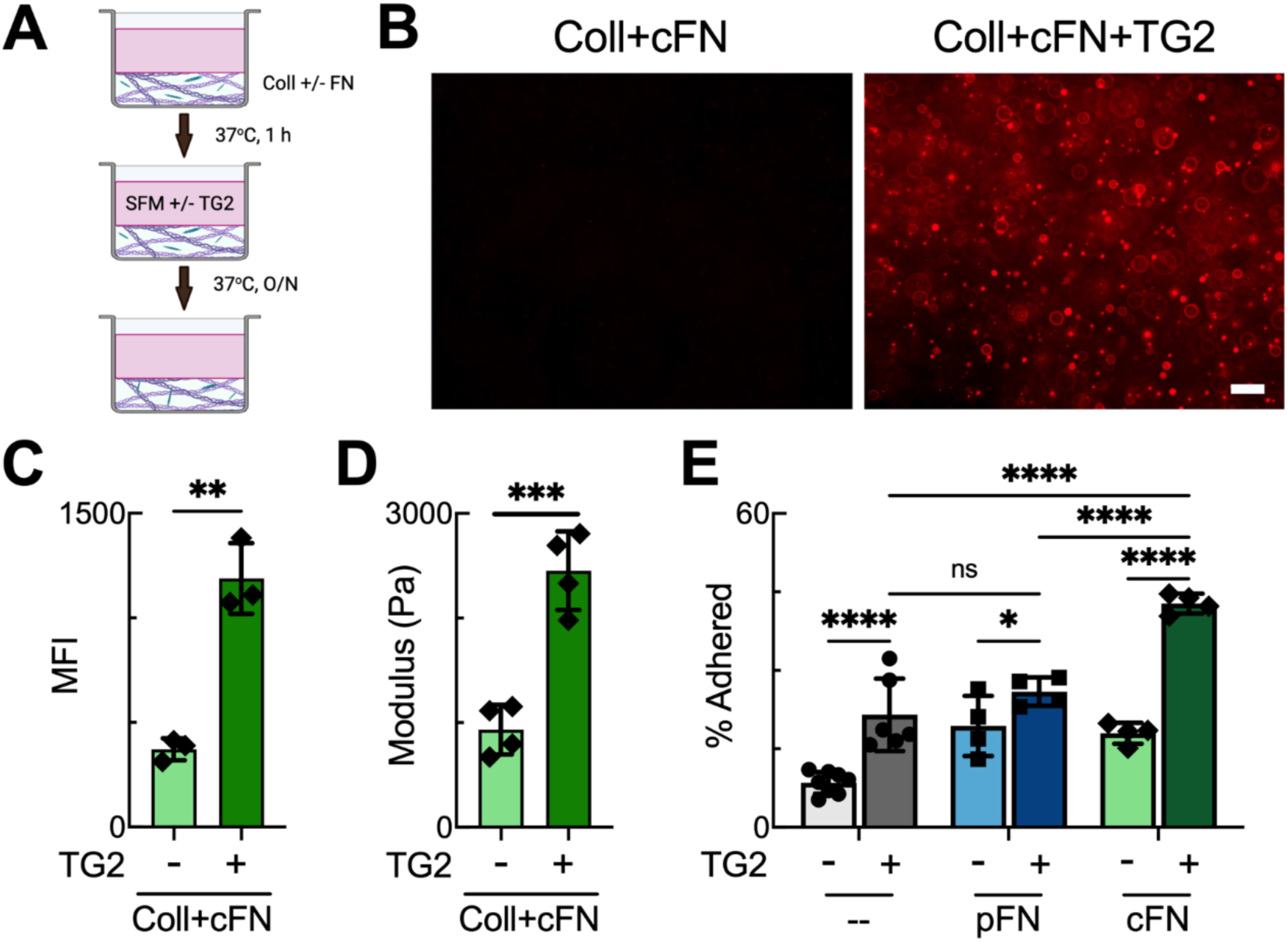
Cross-linking gels with transglutaminase-2 increases cell adhesion. **A,** Schematic depiction of the preparation of TG2 cross-linked gels. **B,** Representative images of N-εγGK-immunostained gels formulated with coll+cFN+/-TG2. Scale bar = 50 μm. **C,** Quantification of N-εγGK MFI for samples in **(B)**. Each dot represents the average across at least 3 fields of view for an individual gel. ** indicates *p*<0.01 by unpaired t-test. **D,** TG2 cross-linking increases the elastic modulus. Each dot represents an individual gel. *** indicates p<0.001 by unpaired t-test. **E,** Quantification of OVCAR4 cells attached on various gel formulations. Each dot represents an individual gel, * indicates p<0.05, **** indicates p<0.0001 by mixed-effects model and Tukey. Data for gels without TG2 are the same as in Figure 2F and the statistical comparison across – TG2 conditions was not included here.

### Integrin engagement and fiber architecture vary between gel formulations

Given that TG2 cross-linked colI/cFN constructs supported significantly increased attachment of HGSOC cells compared to other TG2 cross-linked constructs, we next investigated potential mechanisms for this response. Since integrins are the largest class of receptors mediating cell-ECM adhesion [28, 29] and play critical roles in cancer progression and metastasis [30], we first explored potential changes in integrin engagement upon seeding cells on different hydrogels.

Immunostaining was performed to compare the level of activated β_1_ integrin, a component of many integrin heterodimers known to bind to collagen or fibronectin, in HGSOC cells on coll+TG2 or coll/cFN+TG2 constructs (Fig. 5A,B; Fig. S5A). A significantly elevated level of β_1_ activation was observed in all three HGSOC cell lines after 30 minutes of adhesion on coll/cFN+TG2 compared to coll+TG2. To further investigate the involvement of β_1_ integrin binding, we considered the alpha integrin subunits that provide specificity for the interaction with fibronectin vs. collagen [28]. HGSOC cells were treated with irigenin, a hydroxyisoflavone that blocks the interaction between the EDA domain of cFN with α_9_β_1_ and α_4_β_1_ integrins, or volociximab, a function-blocking antibody specific for integrin α_5_β_1_ and the Arg-Gly-Asp (RGD) motif of fibronectin, and assessed for cell adhesion. Both irigenin and volociximab treatment significantly reduced the adhesion of HGSOC cells to the coll/cFN+TG2 construct (Fig. 5C,D; Fig. S5B,S5C). Interestingly, volociximab appeared to be more potent in inhibiting HGSOC cell adhesion, suggesting that the α_5_β_1_/RGD interaction plays an essential role in mediating HGSOC cell adhesion to col/cFN+TG2.

**Figure 5.**
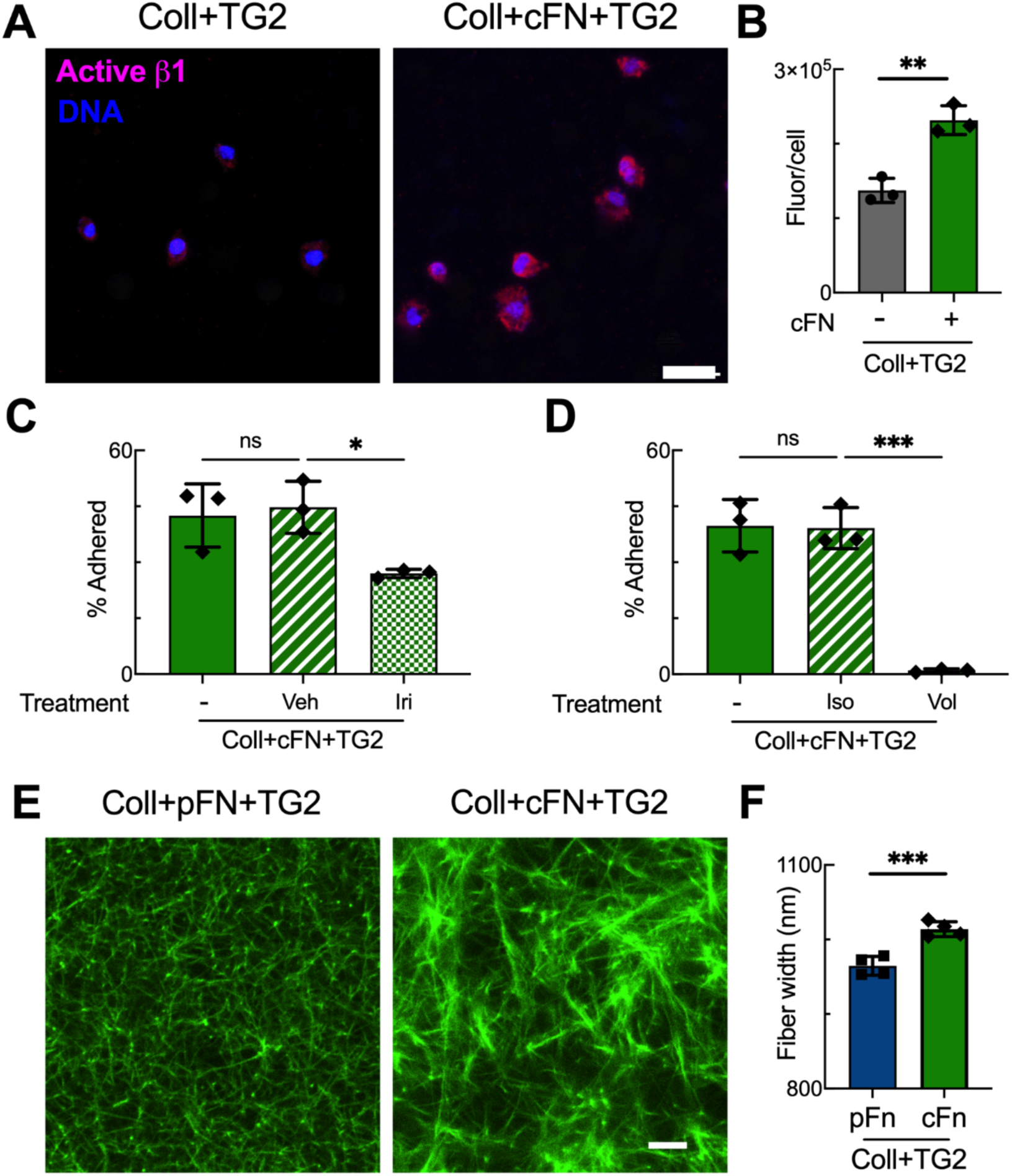
Integrin engagement and fiber architecture vary between different gel formulations. **A,** Representative images of activated β_1_ integrin immunostaining of OVCAR4 cells adhered to coll+TG2 or coll+cFN+TG2 gels. Scale bar = 25 μm. **B,** Quantification of the total fluorescence/cell of activated β_1_ integrin for conditions in **(A)**. Each dot represents the average across at least 3 fields of view for an individual gel, ** indicates *p*<0.01 by unpaired t-test. **C, D,** Quantification of the adhesion of OVCAR4 cells on coll+cFN+TG2 gels upon different treatments. –, no treatment control; Veh, DMSO control; Iri, irigenin; Iso, isotype control; Vol, volociximab. Each dot represents an individual gel, * indicates p<0.05, *** indicates p<0.001 by Tukey. **E,** Representative confocal images of CNA35-EGFP labeled coll+pFN+TG2 and coll+cFN+TG2 gels. Scale bar = 20 μm. **F,** CT-FIRE analysis of mean fiber width in the gels in **(E).** Each dot represents an individual gel, *** indicates p<0.001 by unpaired t-test.

The dependency of HGSOC cell adhesion on the α_5_β_1_/RGD interaction does not fully explain why significantly more cells attached to coll/cFN+TG2 vs. coll/pFN+TG2, as pFN also contains the RGD sequence and the N-terminal collagen binding sites [31]. Therefore, we considered other potential differences that might result from inclusion of cFN versus pFN. cFN has been shown to form fibrils faster than pFN [32], suggesting that it may incorporate differently into the collagen fibers that are forming prior to TG2 cross-linking. CT-FIRE analysis of the CNA35-EGFP labeled collagen fibers revealed large differences in the collagen fiber structure between the coll/cFN+TG2 and the coll/pFN+TG2 constructs (Fig. 5E,F). While the colI/pFN+TG2 construct had numerous short, thin fibers, the coll/cFN+TG2 gel had a distribution of thinner fibers as well as thick fiber bundles. We quantified the fiber widths and found the mean width was significantly increased in the coll/cFN+TG2 construct compared with the colI/pFN+TG2 construct.

### HGSOC cell adhesion is impacted by the source of fibronectin and fiber width

Combined, our results suggest that HGSOC cell adhesion was impacted by fibronectin identity and fiber width in the ECM constructs. However, these two factors appear to interact, whereby the form of fibronectin impacts fiber size (Fig. 5F). Therefore, we sought to identify construct formulations where 1) the fiber width was constant while the form of fibronectin varied, and 2) where the form of fibronectin was constant while the fiber width varied. CNA35-EGFP staining demonstrated that mean fiber width was similar for coll and coll/pFN (Fig. 6A) but was significantly smaller for coll/cFN (Fig. 6B). Prior research has demonstrated that changing the temperature during gelation can impact collagen fiber width [6]; specifically, a slower polymerization at 4°C results in thicker fibers. We tested this protocol with coll/cFN and found that gels with a 1.5 hour incubation at 4°C had a similar mean fiber width as coll and coll/pFN (Fig. S6). We next evaluated cell adhesion across this set of constructs and found that, when fiber width was held constant, incorporation of cFN resulted in the highest number of cells attached (Fig. 6A, Fig. S7A).

**Figure 6.**
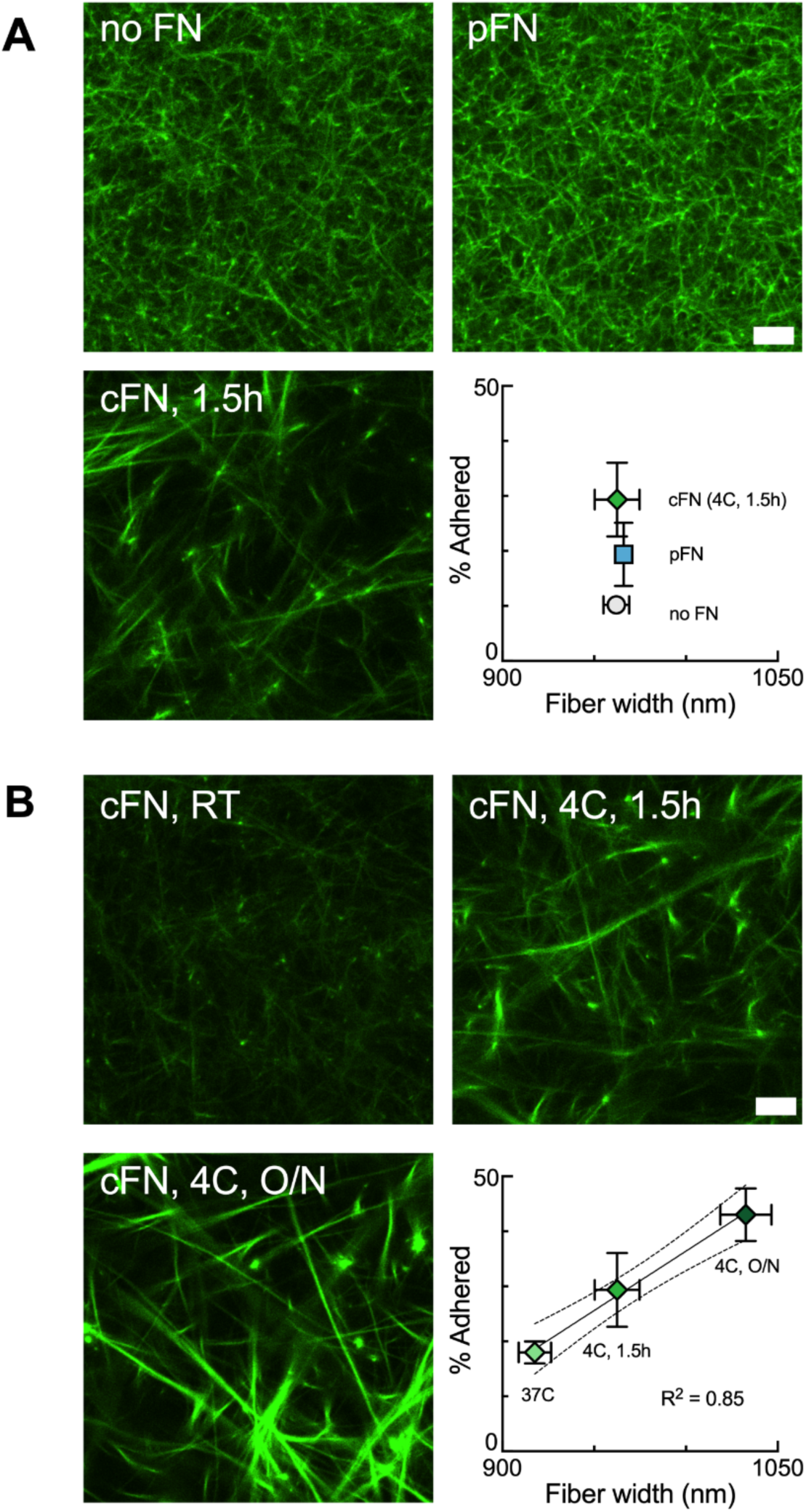
HGSOC cell adhesion is impacted by both fibronectin type and fiber width. **A,** Representative confocal images of CNA35-EGFP stained coll gels with different fibronectin composition but similar mean fiber width. Quantification of mean fiber width and OVCAR4 cell adhesion for the different gel conditions are represented as average ± standard deviation, n = 3-4 gels per measurement. Scale bar = 10 μm. **B,** Representative confocal images of CNA35-EGFP stained coll+cFN gels that were gelled with different conditions to result in a range of fiber widths. Quantification of fiber width and OVCAR4 cell adhesion for the different gel conditions are represented as average ± standard deviation, n = 3-4 gels per measurement. Scale bar = 10 μm.

To examine the impact of fiber width, we examined constructs composed of coll/cFN that were gelled either at 37°C or first incubated for 1.5 hours or overnight at 4°C. As predicted, more time at a lower temperature resulted in even thicker fibers; in fact, the fibers in gels formed overnight were comparable in size to those formed with the addition of TG2 (Fig. 5). When we examined adhesion on gels with varied fiber widths, we observed a strong positive correlation between collagen fiber width and the percentage of HGSOC cells adhered to these constructs (R^2^ = 0.85 for OVCAR4, 0.6 for OVCAR3, and 0.72 for OVCAR8; Fig. 6B, Fig. S7B). In summary, these results indicate that fibronectin identity and fiber width in the ECM constructs act as two independent factors affecting HGSOC cell adhesion.

## Discussion

In the present study, we identified several major changes in the composition and structure of the ECM in HGSOC omental metastases relative to the healthy omentum. Using colIagen I, FN and TG2, we generated fibrillar ECM constructs that recapitulated these key changes and assessed their effects on HGSOC cell adhesion. TG2-mediated cross-linking of colI/cFN hydrogels promoted the adhesion of HGSOC cells through means that were ultimately found to be dependent upon both ligand identity and fiber thickness. HGSOC attachment increased with fiber thickness; however, the composition of these fibers also mediated the extent of adhesion, wherein fibers created by TG2 cross-linking of coll gels containing cFN were more adhesive than those containing pFN. Analysis of gel architecture suggests that the inclusion of cFN vs. pFN impacts collagen fiber formation, resulting in differential cellular adhesion.

To identify changes in the ECM that occur in HGSOC, we analyzed normal omentum and HGSOC omental metastases and observed increases in both collagen I and cFN levels, as well as thickened fibers (Fig. 1, 2). These results are consistent with prior studies [18, 19], and similar compositional changes have been linked with aggressive tumor progression and poor prognosis [33, 34]. Thickening of ECM fibers and other changes to ECM architecture (e.g., alignment) have been observed in primary ovarian tumors [35–37], as well as in other tumor types where aligned or thicker fibers correlate with poor patient survival [4, 38]. The present study observed an overall thickening of the collagen fibers within the metastatic tumor in the omentum. In addition, our lab has previously detected increased fiber density, alignment and width in the collagen underneath the mesothelial layer in tumor-free regions of Stage III/IV omenta [22]. Such changes in ECM fiber architecture may result from protein cross-linking [39], which led us to focus on TG2, an enzyme that is elevated in ovarian cancer and associated with poor survival [40]. While cellular activities of TG2 include promoting epithelial–mesenchymal transition (EMT) and tumor cell survival [40–42], the impact of TG2 on ECM fibers has been relatively understudied. In patients with colorectal cancer, high TG2 expression in the tumor was correlated with thicker collagen fibers and was associated with poor outcomes [43]. We found that TG2 was overexpressed in human HGSOC omental metastases, enzymatically active, and located in the extracellular environment. Therefore, we sought to incorporate the relative contributions of increased collagen, FN, and TG2 enzymatic activity in our *in vitro* ECM models and examine their effects on fiber structure and cell adhesion.

Both naturally-derived materials and synthetic approaches have been used to construct ECM-mimetic scaffolds to study tumor metastasis [1]. For example, decellularized matrices have been prepared from many tissues, including mammary glands, ovary, omentum and peritoneum [44–47]. However, decellularized matrices do not allow the contribution of individual ECM components to be isolated and are subject to challenges with reproducibility. At the other end of the spectrum are entirely synthetic hydrogels that allow for exquisite control over scaffold composition and are highly reproducible but typically do not replicate the architectural elements of the ECM. For example, polyethylene glycol (PEG) gels have been functionalized with integrin-binding peptides from collagen and fibronectin in order to examine chemotherapy resistance in a model of omental HGSOC metastases [48]. Here, we opted for an intermediate option, using naturally derived, individual components of the ECM that reproduce native, fibrillar ECM architecture and whose composition can be tailored based on the quantitative analysis of patient samples. While increased collagen has been suggested as a key reason that tumor cells metastasize to the omentum [49, 50], we found that a low percentage of HGSOC cells adhered to colI gels. We next considered the impact of fibronectin and incorporated pFN into colI gels, as other cell types exhibit different behaviors when coll hydrogels are blended with pFN. For example, incorporation of pFN into colI hydrogels facilitated focal adhesion formation and cell spreading [51, 52], although this work did not use tumor cells. Our studies found increased attachment when pFN was incorporated into colI gels for only one of the three HGSOC cell lines examined. We next examined whether these findings were dependent upon the type of FN. cFN is also known to promote cell adhesion and migration but has a distinct molecular structure and functional domains. Whether cFN and pFN are equally active in promoting cell adhesion and spreading appears to be dependent on the cell types examined [53, 54]. For example, ovarian surface epithelial cells and SKOV-3 attached and spread faster on cFN, whereas OVCAR3 cells displayed similar attachment between cFN and pFN [54]. We found that all three HGSOC cell lines showed similar attachment between colI/cFN and colI/pFN gels, with a significant increase relative to coll observed for only one cell line. When gels were formed using a standard gelation protocol, we did not observe differences between the isoforms. However, we did observe increased adhesion on cFN relative to pFN when the fiber structure was manipulated to achieve equivalent fiber thickness through gelation temperature or when TG2 was used to cross-link the network.

It has been increasingly recognized that the presence of ECM fibers is crucial for cancer progression. For example, by incorporating collagen fibers into a gelatin-methacrylate network *in vitro*, our group illustrated that the presence of collagen fibers was required for breast cancer cells to invade the surrounding matrix [3]. Moreover, prior studies have demonstrated that features such as fiber alignment, thickness, length and density, significantly influence tumor behavior and correlate with clinical outcomes in other cancers. For example, in breast cancer, radially aligned collagen fibers facilitate local cancer cell invasion [4]; while in pancreatic ductal carcinoma, thicker collagen fibers adjacent to lesions are linked to shorter survival [55]. In HGSOC, collagen fibers in the stroma of the primary tumor display increased alignment and wavy morphology, which *in vitro* studies suggest may enhance ovarian cancer cell motility [35, 56]. Several methods have been established to manipulate collagen fibril diameter *in vitro* by controlling pH, collagen concentration and temperature of gelation [6, 57]. In contrast, ECM fiber thickening *in vivo* results from new protein deposition, enzymatic cross-linking and fiber bundling [58]. Here, we treated colI, colI/pFN, and colI/cFN constructs with TG2 to induce cross-linking, which was confirmed by the presence of the N-epsilon gamma-glutamyl lysine cross-links and an increase in bulk stiffness. While TG2-mediated cross-linking of coll and coll/pFN gels increased cell adhesion for a subset of conditions, the TG2 cross-linked colI/cFN gel had the highest level of HGSOC cell attachment for all HGSOC cells examined. Thus, in the presence of TG2, the type of FN used in the biomaterial construct affects the adhesion of HGSOC cells.

We next examined the mechanisms responsible for the marked increase in adhesion on TG2-cross-linked coll/cFN constructs. A key feature of adhesion complexes is the integrin receptor, which provides specificity for the cell-ECM interaction through the α and β subunits involved. Collagen and cFN both interact with the β_1_ subunit but utilize different α subunit partners; specifically, collagen binds to α_1_β_1_, α_2_β_1,_ α_10_β_1,_ and α_11_β_1_ while cFN interacts with α_5_β_1_, α_4_β_1_ and α_9_β_1_ [59, 60]. The increase in activated β_1_ integrin on TG2 cross-linked coll/cFN gels reported here led us to hypothesize that cFN-specific interactions were responsible. Using inhibitors against the RGD/α_5_β_1_ and EDA/α_4_β_1_ or α_9_β_1_ interaction, we determined that the RGD motif was essential for the increase in adhesion. However, the central role for RGD leads to a conundrum as pFN contains the necessary RGD domain and N-terminal collagen binding sites [61], but did not consistently lead to increased tumor cell adhesion. Our recent work demonstrated that collagen fibers support the formation of nascent adhesions which ultimately lead to more mature adhesion complexes in HGSOC cells [22]. Therefore, one potential explanation for this cFN-specific increase on TG2 cross-linked gels is the observation that TG2 cross-linked coll/cFN gels had significantly thicker fibers than coll/pFN gels. Many studies of collagen/fibronectin interactions during fiber formation use the plasma form of fibronectin. However, it has been reported that cFN is assembled more efficiently into the ECM compared to pFN, potentially due to the enhanced self-polymerization when the EDA/EDB domains are present [62, 63]. Recent work has demonstrated that TG2 has a higher affinity for relaxed fibronectin fibers [64]; while this work was conducted with pFN, it suggests that the differences in conformation resulting from the additional domains in cFN may impact the coll/coll vs. coll/FN cross-links formed in the different gels. An alternative explanation is that soluble pFN adopts a closed form with the RGD motif unavailable for cell binding [61] while the EDA domain present in cFN may induce conformational changes that improve the accessibility of the RGD site for engagement by cell surface receptors [61, 65, 66]. In addition, cFN contains a non-homologous variable (V) region (also known as the type III connecting segment (IIICS) domain), which is lacking in one subunit of the pFN dimer [13]. Evidence suggests that the IIICS domain might be required for the EDA-dependent enhancement of cell adhesion to FN [66]. While we did not observe significant differences in adhesion between cFN and pFN when simply mixed, it is possible that these conformational differences are more pronounced following TG2-mediated cross-linking.

## Conclusion

Cell adhesion is a fundamental requirement for anchorage-dependent cells to survive and is an essential step during transcoelomic spread, where cancer cells must attach to peritoneal tissues to initiate a new metastatic site. Our prior results demonstrated that the density of ECM fibers impacted HGSOC cell adhesion [22]; here, we demonstrate that the TG2-colI-cFN triad plays a critical role in modulating ECM fiber width and HGSOC cell attachment. Since these three components of the ECM are overexpressed in many cancers [60, 67], we propose that our constructs will be useful to study the impact of ECM architecture on a variety of cellular processes at play during tumor progression and metastasis. Importantly, our results demonstrate that the more commonly used pFN does not produce the same effects as cFN when cross-linked by TG2 and suggest that this difference is a result of changes in fiber structure.

## Data Availability

All the datasets generated during and analyzed during the current study are available from the corresponding authors upon reasonable request.

## Funding and Acknowledgements

This work was supported by National Institutes for Health research grants R01CA232517 (PKK, KSM) and R01CA290693 (PKK), and the University of Wisconsin Graduate School (PKK). The authors gratefully acknowledge the patients who consented for their tissue and ascites samples to be collected and the University of Wisconsin Carbone Cancer Center BioBank, supported by P30 CA014520. The authors thank the University of Wisconsin Translational Research Initiatives in Pathology (TRIP) Laboratory, supported by the UW Department of Pathology and Laboratory Medicine, UWCCC (P30 CA014520) and the Office of the NIH Director (S10 OD023526) for use of its facilities and services. We thank the University of Wisconsin Optical Imaging Core for use of the Nikon AXR LSCM, supported by the NIH Office of the NIH Director (S10 OD034394-0). The authors gratefully acknowledge use of facilities and instrumentation in the UW-Madison Wisconsin Center for Nanoscale Technology. The Center (wcnt.wisc.edu) is partially supported by the Wisconsin Materials Research Science and Engineering Center (NSF DMR-2309000) and the University of Wisconsin-Madison. The funders had no role in study design, data collection, or analysis.

## Notes

### Competing Interest Statement

The authors have declared no competing interest.

## References

[1] H.M. Micek, M.R. Visetsouk, K.S. Masters, P.K. Kreeger, Engineering the Extracellular Matrix to Model the Evolving Tumor Microenvironment, iScience 23(11) (2020) 101742.

[2] J.C. Ashworth, T.R. Cox, The importance of 3D fibre architecture in cancer and implications for biomaterial model design, Nat Rev Cancer 24(7) (2024) 461–479.

[3] A.J. Berger, K.M. Linsmeier, P.K. Kreeger, K.S. Masters, Decoupling the effects of stiffness and fiber density on cellular behaviors via an interpenetrating network of gelatin-methacrylate and collagen, Biomaterials 141 (2017) 125–135.

[4] P.P. Provenzano, K.W. Eliceiri, J.M. Campbell, D.R. Inman, J.G. White, P.J. Keely, Collagen reorganization at the tumor-stromal interface facilitates local invasion, BMC Med 4(1) (2006) 38.

[5] A. Naba, Mechanisms of assembly and remodelling of the extracellular matrix, Nat Rev Mol Cell Biol 25(11) (2024) 865–885.

[6] B.R. Seo, X. Chen, L. Ling, Y.H. Song, A.A. Shimpi, S. Choi, J. Gonzalez, J. Sapudom, K. Wang, R.C. Andresen Eguiluz, D. Gourdon, V.B. Shenoy, C. Fischbach, Collagen microarchitecture mechanically controls myofibroblast differentiation, Proc Natl Acad Sci U S A 117(21) (2020) 11387–11398.

[7] D. Ceballos, X. Navarro, N. Dubey, G. Wendelschafer-Crabb, W.R. Kennedy, R.T. Tranquillo, Magnetically aligned collagen gel filling a collagen nerve guide improves peripheral nerve regeneration, Exp Neurol 158(2) (1999) 290–300.

[8] C. Guo, L.J. Kaufman, Flow and magnetic field induced collagen alignment, Biomaterials 28(6) (2007) 1105–14.

[9] P. Lee, R. Lin, J. Moon, L.P. Lee, Microfluidic alignment of collagen fibers for in vitro cell culture, Biomed Microdevices 8(1) (2006) 35–41.

[10] K.M. Riching, B.L. Cox, M.R. Salick, C. Pehlke, A.S. Riching, S.M. Ponik, B.R. Bass, W.C. Crone, Y. Jiang, A.M. Weaver, K.W. Eliceiri, P.J. Keely, 3D collagen alignment limits protrusions to enhance breast cancer cell persistence, Biophys J 107(11) (2014) 2546–58.

[11] A. Ray, O. Lee, Z. Win, R.M. Edwards, P.W. Alford, D.H. Kim, P.P. Provenzano, Anisotropic forces from spatially constrained focal adhesions mediate contact guidance directed cell migration, Nat Commun 8 (2017) 14923.

[12] H. Jiang, M. Zheng, X. Liu, S. Zhang, X. Wang, Y. Chen, M. Hou, J. Zhu, Feasibility Study of Tissue Transglutaminase for Self-Catalytic Cross-Linking of Self-Assembled Collagen Fibril Hydrogel and Its Promising Application in Wound Healing Promotion, ACS Omega 4(7) (2019) 12606–12615.

[13] C.J. Dalton, C.A. Lemmon, Fibronectin: Molecular Structure, Fibrillar Structure and Mechanochemical Signaling, Cells 10(9) (2021).

[14] K.E. Kadler, A. Hill, E.G. Canty-Laird, Collagen fibrillogenesis: fibronectin, integrins, and minor collagens as organizers and nucleators, Curr Opin Cell Biol 20(5) (2008) 495–501.

[15] C.A. Sevilla, D. Dalecki, D.C. Hocking, Regional fibronectin and collagen fibril co-assembly directs cell proliferation and microtissue morphology, PLoS One 8(10) (2013) e77316.

[16] J.H. Pincus, H. Waelsch, The specificity of transglutaminase. II. Structural requirements of the amine substrate, Arch Biochem Biophys 126(1) (1968) 44–52.

[17] M.V. Nurminskaya, A.M. Belkin, Cellular functions of tissue transglutaminase, Int Rev Cell Mol Biol 294 (2012) 1–97.

[18] K.C. Fogg, C.M. Renner, H. Christian, A. Walker, L. Marty-Santos, A. Khan, W.R. Olson, C. Parent, A. O’Shea, D.M. Wellik, P.S. Weisman, P.K. Kreeger, Ovarian Cells Have Increased Proliferation in Response to Heparin-Binding Epidermal Growth Factor as Collagen Density Increases, Tissue Eng Part A 26(13-14) (2020) 747–758.

[19] O.M.T. Pearce, R.M. Delaine-Smith, E. Maniati, S. Nichols, J. Wang, S. Bohm, V. Rajeeve, D. Ullah, P. Chakravarty, R.R. Jones, A. Montfort, T. Dowe, J. Gribben, J.L. Jones, H.M. Kocher, J.S. Serody, B.G. Vincent, J. Connelly, J.D. Brenton, C. Chelala, P.R. Cutillas, M. Lockley, C. Bessant, M.M. Knight, F.R. Balkwill, Deconstruction of a Metastatic Tumor Microenvironment Reveals a Common Matrix Response in Human Cancers, Cancer Discov 8(3) (2018) 304–319.

[20] J.S. Bredfeldt, Y. Liu, C.A. Pehlke, M.W. Conklin, J.M. Szulczewski, D.R. Inman, P.J. Keely, R.D. Nowak, T.R. Mackie, K.W. Eliceiri, Computational segmentation of collagen fibers from second-harmonic generation images of breast cancer, J Biomed Opt 19(1) (2014) 16007.

[21] S.J. Aper, A.C. van Spreeuwel, M.C. van Turnhout, A.J. van der Linden, P.A. Pieters, N.L. van der Zon, S.L. de la Rambelje, C.V. Bouten, M. Merkx, Colorful protein-based fluorescent probes for collagen imaging, PLoS One 9(12) (2014) e114983.

[22] A. Abbaspour, A.L. Martinez Cavazos, R. Patel, N. Yang, S.M. McGregor, E.G. Brooks, K.S. Masters, P.K. Kreeger, Collagen Fiber Density Observed in Metastatic Ovarian Cancer Promotes Tumor Cell Adhesion, Acta Biomater. (accepted).

[23] M.C. Erat, B. Sladek, I.D. Campbell, I. Vakonakis, Structural analysis of collagen type I interactions with human fibronectin reveals a cooperative binding mode, J Biol Chem 288(24) (2013) 17441–50.

[24] Y. Sun, A.J. Hamlin, J.E. Schwarzbauer, Fibronectin matrix assembly at a glance, J Cell Sci 138(6) (2025).

[25] S.E. Bruce, T.J. Peters, The subcellular localization of transglutaminase in normal liver and in glucagon-treated and partial hepatectomized rats, Biosci Rep 3(12) (1983) 1085–90.

[26] H. Tatsukawa, K. Hitomi, Role of Transglutaminase 2 in Cell Death, Survival, and Fibrosis, Cells 10(7) (2021).

[27] K.R. Levental, H. Yu, L. Kass, J.N. Lakins, M. Egeblad, J.T. Erler, S.F. Fong, K. Csiszar, A. Giaccia, W. Weninger, M. Yamauchi, D.L. Gasser, V.M. Weaver, Matrix crosslinking forces tumor progression by enhancing integrin signaling, Cell 139(5) (2009) 891–906.

[28] I.D. Campbell, M.J. Humphries, Integrin structure, activation, and interactions, Cold Spring Harb Perspect Biol 3(3) (2011).

[29] M.R. Chastney, J.R.W. Conway, J. Ivaska, Integrin adhesion complexes, Curr Biol 31(10) (2021) R536–R542.

[30] M.R. Chastney, J. Kaivola, V.M. Leppänen, J. Ivaska, The role and regulation of integrins in cell migration and invasion, Nat Rev Mol Cell Biol 26(2) (2025) 147–167.

[31] J. Cho, D.F. Mosher, Role of fibronectin assembly in platelet thrombus formation, J Thromb Haemost 4(7) (2006) 1461–9.

[32] J.L. Sechler, Y. Takada, J.E. Schwarzbauer, Altered rate of fibronectin matrix assembly by deletion of the first type III repeats, J Cell Biol 134(2) (1996) 573–83.

[33] K.A. Kujawa, E. Zembala-Nozynska, A.J. Cortez, T. Kujawa, J. Kupryjanczyk, K.M. Lisowska, Fibronectin and Periostin as Prognostic Markers in Ovarian Cancer, Cells 9(1) (2020).

[34] X. Xiao, F. Long, S. Yu, W. Wu, D. Nie, X. Ren, W. Li, X. Wang, L. Yu, P. Wang, G. Wang, Col1A1 as a new decoder of clinical features and immune microenvironment in ovarian cancer, Front Immunol 15 (2024) 1496090.

[35] B. Wen, K.R. Campbell, K. Tilbury, O. Nadiarnykh, M.A. Brewer, M. Patankar, V. Singh, K.W. Eliceiri, P.J. Campagnola, 3D texture analysis for classification of second harmonic generation images of human ovarian cancer, Sci Rep 6 (2016) 35734.

[36] K.R. Campbell, P.J. Campagnola, Assessing local stromal alterations in human ovarian cancer subtypes via second harmonic generation microscopy and analysis, J Biomed Opt 22(11) (2017) 1–7.

[37] M. Sarwar, P.H. Sykes, K. Chitcholtan, J.J. Evans, Collagen I dysregulation is pivotal for ovarian cancer progression, Tissue Cell 74 (2022) 101704.

[38] C.R. Drifka, A.G. Loeffler, K. Mathewson, A. Keikhosravi, J.C. Eickhoff, Y. Liu, S.M. Weber, W.J. Kao, K.W. Eliceiri, Highly aligned stromal collagen is a negative prognostic factor following pancreatic ductal adenocarcinoma resection, Oncotarget 7(46) (2016) 76197–76213.

[39] Z. Wang, M. Griffin, TG2, a novel extracellular protein with multiple functions, Amino Acids 42(2-3) (2012) 939–49.

[40] J.Y. Hwang, L.S. Mangala, J.Y. Fok, Y.G. Lin, W.M. Merritt, W.A. Spannuth, A.M. Nick, D.J. Fiterman, P.E. Vivas-Mejia, M.T. Deavers, R.L. Coleman, G. Lopez-Berestein, K. Mehta, A.K. Sood, Clinical and biological significance of tissue transglutaminase in ovarian carcinoma, Cancer Res 68(14) (2008) 5849–58.

[41] L. Cao, D.N. Petrusca, M. Satpathy, H. Nakshatri, I. Petrache, D. Matei, Tissue transglutaminase protects epithelial ovarian cancer cells from cisplatin-induced apoptosis by promoting cell survival signaling, Carcinogenesis 29(10) (2008) 1893–900.

[42] M. Shao, L. Cao, C. Shen, M. Satpathy, B. Chelladurai, R.M. Bigsby, H. Nakshatri, D. Matei, Epithelial-to-mesenchymal transition and ovarian tumor progression induced by tissue transglutaminase, Cancer Res 69(24) (2009) 9192–201.

[43] R. Delaine-Smith, N. Wright, C. Hanley, R. Hanwell, R. Bhome, M. Bullock, C. Drifka, K. Eliceiri, G. Thomas, M. Knight, A. Mirnezami, N. Peake, Transglutaminase-2 Mediates the Biomechanical Properties of the Colorectal Cancer Tissue Microenvironment that Contribute to Disease Progression, Cancers (Basel) 11(5) (2019).

[44] L. Varinelli, M. Guaglio, S. Brich, S. Zanutto, A. Belfiore, F. Zanardi, F. Iannelli, A. Oldani, E. Costa, M. Chighizola, E. Lorenc, S.P. Minardi, S. Fortuzzi, M. Filugelli, G. Garzone, F. Pisati, M. Vecchi, G. Pruneri, S. Kusamura, D. Baratti, L. Cattaneo, D. Parazzoli, A. Podestà, M. Milione, M. Deraco, M.A. Pierotti, M. Gariboldi, Decellularized extracellular matrix as scaffold for cancer organoid cultures of colorectal peritoneal metastases, J Mol Cell Biol 14(11) (2023).

[45] A.L. Wishart, S.J. Conner, J.R. Guarin, J.P. Fatherree, Y. Peng, R.A. McGinn, R. Crews, S.P. Naber, M. Hunter, A.S. Greenberg, M.J. Oudin, Decellularized extracellular matrix scaffolds identify full-length collagen VI as a driver of breast cancer cell invasion in obesity and metastasis, Sci Adv 6(43) (2020).

[46] T. Wu, Y.Y. Gao, X.N. Tang, J.J. Zhang, S.X. Wang, Construction of Artificial Ovaries with Decellularized Porcine Scaffold and Its Elicited Immune Response after Xenotransplantation in Mice, J Funct Biomater 13(4) (2022).

[47] A. Emet, E. Ozdemir, D.U. Cetinkaya, Kilic, E., R. Hashemihesar, A.C.S. Yuruker, E. Turhan, Effect of a decellularized omentum scaffold with combination of mesenchymal stem cells and platelet-rich plasma on healing of critical-sized bone defect: a rat model., Appl. Sci., 2021, p. 10900.

[48] E.A. Brooks, M.F. Gencoglu, D.C. Corbett, K.R. Stevens, S.R. Peyton, An omentum-inspired 3D PEG hydrogel for identifying ECM-drivers of drug resistant ovarian cancer, APL Bioeng 3(2) (2019) 026106.

[49] E.W. Sorensen, S.A. Gerber, A.L. Sedlacek, V.Y. Rybalko, W.M. Chan, E.M. Lord, Omental immune aggregates and tumor metastasis within the peritoneal cavity, Immunol Res 45(2-3) (2009) 185–94.

[50] Y.L. Huang, C.Y. Liang, D. Ritz, R. Coelho, D. Septiadi, M. Estermann, C. Cumin, N. Rimmer, A. Schötzau, M. Núñez López, A. Fedier, M. Konantz, T. Vlajnic, D. Calabrese, C. Lengerke, L. David, B. Rothen-Rutishauser, F. Jacob, V. Heinzelmann-Schwarz, Collagen-rich omentum is a premetastatic niche for integrin α2-mediated peritoneal metastasis, Elife 9 (2020).

[51] Y. Nakamura, T. Sagara, K. Seki, S. Hirano, T. Nishida, Permissive effect of fibronectin on collagen gel contraction mediated by bovine trabecular meshwork cells, Invest Ophthalmol Vis Sci 44(10) (2003) 4331–6.

[52] Y. Nashchekina, P. Nikonov, N. Prasolov, M. Sulatsky, A. Chabina, A. Nashchekin, The Structural Interactions of Molecular and Fibrillar Collagen Type I with Fibronectin and Its Role in the Regulation of Mesenchymal Stem Cell Morphology and Functional Activity, Int J Mol Sci 23(20) (2022).

[53] K.M. Yamada, D.W. Kennedy, Fibroblast cellular and plasma fibronectins are similar but not identical, J Cell Biol 80(2) (1979) 492–8.

[54] L. Zand, F. Qiang, C.D. Roskelley, P.C. Leung, N. Auersperg, Differential effects of cellular fibronectin and plasma fibronectin on ovarian cancer cell adhesion, migration, and invasion, In Vitro Cell Dev Biol Anim 39(3-4) (2003) 178–82.

[55] H. Laklai, Y.A. Miroshnikova, M.W. Pickup, E.A. Collisson, G.E. Kim, A.S. Barrett, R.C. Hill, J.N. Lakins, D.D. Schlaepfer, J.K. Mouw, V.S. LeBleu, N. Roy, S.V. Novitskiy, J.S. Johansen, V. Poli, R. Kalluri, C.A. Iacobuzio-Donahue, L.D. Wood, M. Hebrok, K. Hansen, H.L. Moses, V.M. Weaver, Genotype tunes pancreatic ductal adenocarcinoma tissue tension to induce matricellular fibrosis and tumor progression, Nat Med 22(5) (2016) 497–505.

[56] S. Alkmin, R. Brodziski, H. Simon, D. Hinton, R.H. Goldsmith, M. Patankar, P.J. Campagnola, Role of Collagen Fiber Morphology on Ovarian Cancer Cell Migration Using Image-Based Models of the Extracellular Matrix, Cancers (Basel) 12(6) (2020).

[57] J. Sapudom, S. Rubner, S. Martin, T. Kurth, S. Riedel, C.T. Mierke, T. Pompe, The phenotype of cancer cell invasion controlled by fibril diameter and pore size of 3D collagen networks, Biomaterials 52 (2015) 367–75.

[58] C. Frantz, K.M. Stewart, V.M. Weaver, The extracellular matrix at a glance, J Cell Sci 123(Pt 24) (2010) 4195–200.

[59] M. Bachmann, S. Kukkurainen, V.P. Hytonen, B. Wehrle-Haller, Cell Adhesion by Integrins, Physiol Rev 99(4) (2019) 1655–1699.

[60] J.W. Rick, A. Chandra, C. Dalle Ore, A.T. Nguyen, G. Yagnik, M.K. Aghi, Fibronectin in malignancy: Cancer-specific alterations, protumoral effects, and therapeutic implications, Semin Oncol 46(3) (2019) 284–290.

[61] E.S. White, F.E. Baralle, A.F. Muro, New insights into form and function of fibronectin splice variants, J Pathol 216(1) (2008) 1–14.

[62] G. Efthymiou, A. Radwanska, A.I. Grapa, S. Beghelli-de la Forest Divonne, D. Grall, S. Schaub, M. Hattab, S. Pisano, M. Poet, D.F. Pisani, L. Counillon, X. Descombes, L. Blanc-Féraud, E. Van Obberghen-Schilling, Fibronectin Extra Domains tune cellular responses and confer topographically distinct features to fibril networks, J Cell Sci 134(4) (2021).

[63] J.L. Guan, J.E. Trevithick, R.O. Hynes, Retroviral expression of alternatively spliced forms of rat fibronectin, J Cell Biol 110(3) (1990) 833–47.

[64] K. Selcuk, A. Leitner, L. Braun, F. Le Blanc, P. Pacak, S. Pot, V. Vogel, Transglutaminase 2 has higher affinity for relaxed than for stretched fibronectin fibers, Matrix Biol 125 (2024) 113–132.

[65] D.J. Leahy, I. Aukhil, H.P. Erickson, 2.0 A crystal structure of a four-domain segment of human fibronectin encompassing the RGD loop and synergy region, Cell 84(1) (1996) 155–64.

[66] R. Manabe, N. Ohe, T. Maeda, T. Fukuda, K. Sekiguchi, Modulation of cell-adhesive activity of fibronectin by the alternatively spliced EDA segment, J Cell Biol 139(1) (1997) 295–307.

[67] N.I. Nissen, M. Karsdal, N. Willumsen, Collagens and Cancer associated fibroblasts in the reactive stroma and its relation to Cancer biology, J Exp Clin Cancer Res 38(1) (2019) 115.

