## Supplemental Tables and Figures for "Fibronectin Composition and Transglutaminase 2 Cross-linking Cooperatively Regulate Ovarian Cancer Cell Adhesion in ECM-Mimetic Constructs"

**Supplementary Table 1.** Patient information for the omental samples.

| Specimen ID | Age | Race, ethnicity (provided by patient) | Clinical Diagnosis |
|---|---|---|---|
| Healthy-1 | 61 | White | N/A |
| Healthy -2 | 63 | White, Hispanic | N/A |
| Healthy -3 | 82 | White | N/A |
| Healthy -4 | 64 | White | N/A |
| Healthy -5 | 84 | White | N/A |
| Healthy -6 | 41 | White | N/A |
| Healthy -7 | 70 | White | N/A |
| Healthy -8 | 88 | White | N/A |
| HGSOC-1 | 58 | White | HGSOC, IIIC |
| HGSOC-2 | 54 | White | HGSOC, IIIB |
| HGSOC-3 | 54 | White | HGSOC, IIIC |
| HGSOC-4 | 74 | White | HGSOC, IIIC |
| HGSOC-5 | 82 | White | HGSOC, IIIC |
| HGSOC-6 | 66 | White | HGSOC, IIIC |
| HGSOC-7 | 76 | White | HGSOC, IIIB |

**Supplementary Table 2.** Patient information for the ascites samples.

| Specimen ID | Age | Race, ethnicity (provided by patient) | Clinical Diagnosis |
|---|---|---|---|
| Benign-1 | 46 | N/A | Serous borderline tumor |
| Benign-2 | 61 | N/A | Fibroma |
| Benign-3 | 56 | N/A | Fibroma |
| Benign-4 | 51 | N/A | Endometrioma |
| HGSOC-1 | 77 | White | HGSOC, IIIC |
| HGSOC-2 | 66 | White | HGSOC, IIIC |
| HGSOC-3 | 79 | White | HGSOC, IIIC |
| HGSOC-4 | 58 | White | HGSOC, IIIC |
| HGSOC-5 | 73 | White | HGSOC, IIIC |
| HGSOC-6 | 71 | Asian | HGSOC, IIIC |
| HGSOC-7 | 69 | American Indian or Alaskin native | HGSOC, IIIC |
| HGSOC-8 | 69 | White | HGSOC, IVB |
| HGSOC-9 | 70 | White | HGSOC, IIIC |
| HGSOC-10 | 50 | N/A | HGSOC, IIIC |
| HGSOC-11 | 61 | N/A | HGSOC, IVA |
| HGSOC-12 | 44 | N/A | HGSOC, IIIC |
| HGSOC-13 | 57 | N/A | HGSOC, IIIC |

**Supplementary Table 3.** Patient information for the metastatic samples analyzed for extracellular TG2.

| Specimen | Age | Race, ethnicity (provided by patient) | Clinical Details |
|---|---|---|---|
| 1 | 48 | White | Stage III, collected at interval debulking (2 cycles) |
| 2 (shown in Fig 3E) | 66 | White | Stage IIIC, collected during primary debulking |

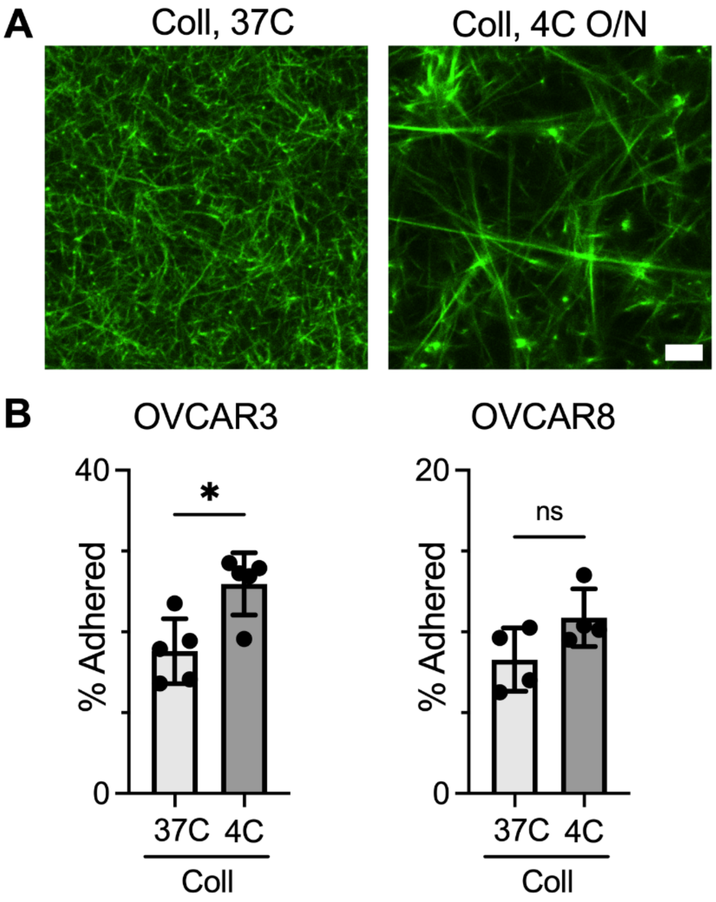


**Supplementary Figure 1.** Thicker fibers in coll gels support increased tumor cell attachment. **A,** Representative CNA35-EGFP stained coll gels formed at 37°C for one hour (37C) or 4°C overnight (4C). Scale bar = 10 μm. **B,** Quantification of OVCAR3 and OVCAR8 cells attached on coll gels. Each dot represents an individual gel, ns indicates $p>0.05$, * indicates $p<0.05$, by unpaired t-test.

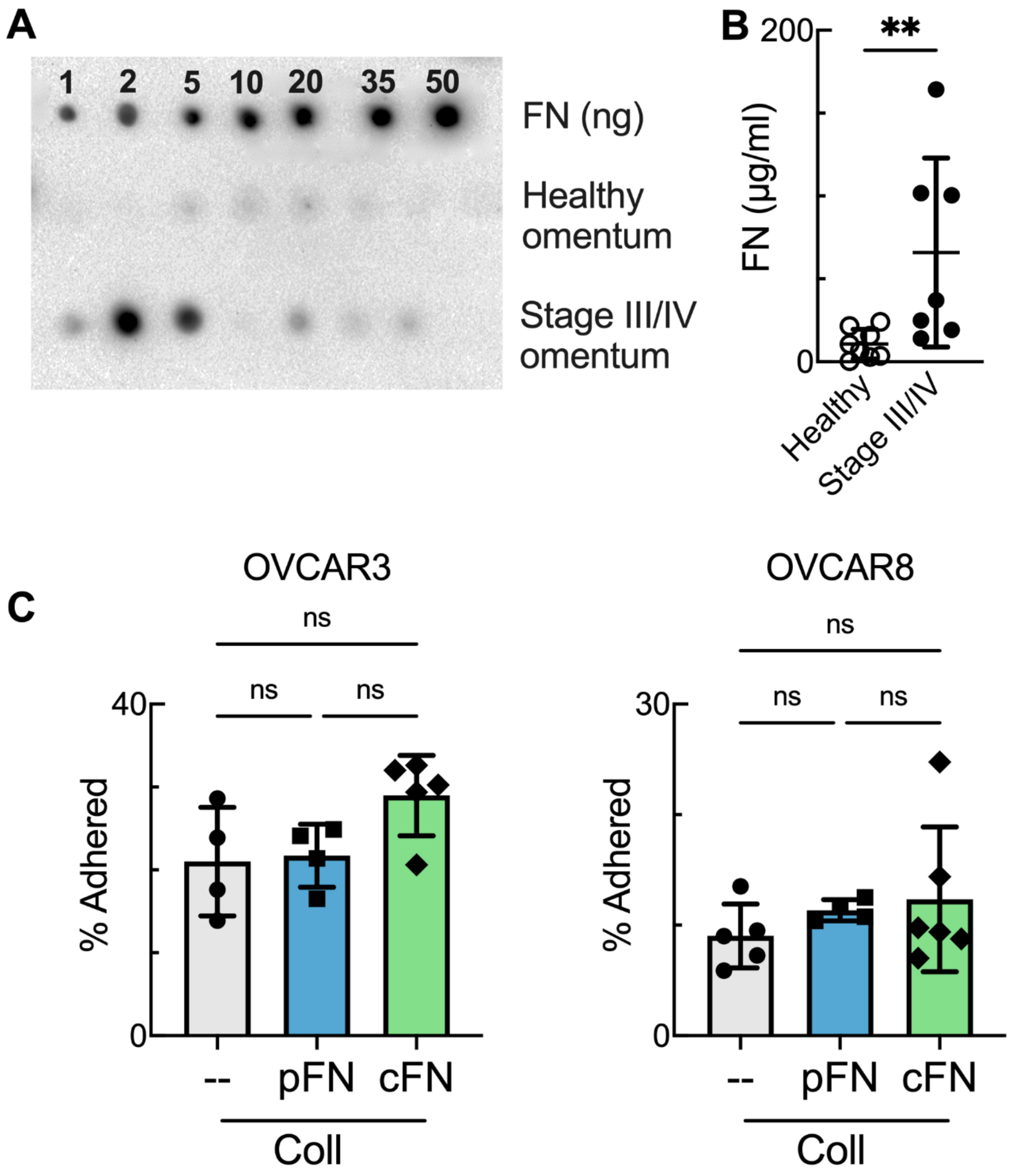


**Supplementary Figure 2.** Fibronectin is increased in human HGSOC omental metastases. **A,** Dot blot probed for fibronectin. Standards are purified fibronectin from human plasma, samples are solubilized protein from sections of healthy omentum or Stage III/IV omental tumors. **B,** Quantification of dot blot. Each dot represents an individual patient, ** indicates $p<0.01$ by unpaired t-test. **C,** Quantification of OVCAR3 and OVCAR8 cells attached on coll, coll+pFN, or coll+cFN. Each dot represents an individual gel. ns indicates $p>0.05$ by ANOVA.

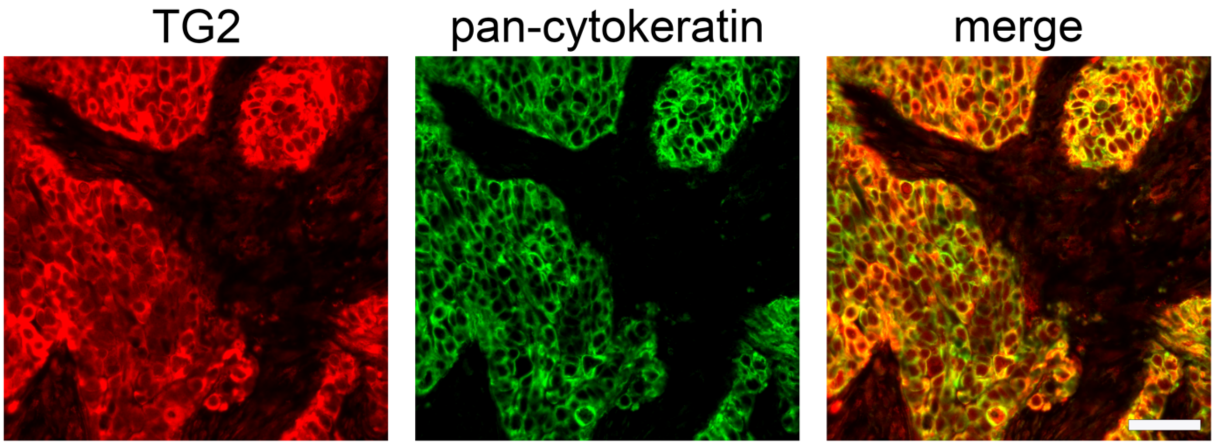


**Supplementary Figure 3.** Transglutaminase is present in both the epithelial and stromal fraction of omental tumors. Representative images of TG2-immunostained human omental samples counterstained for pan-cytokeratin (epithelial marker). Scale bar = 50 µm.

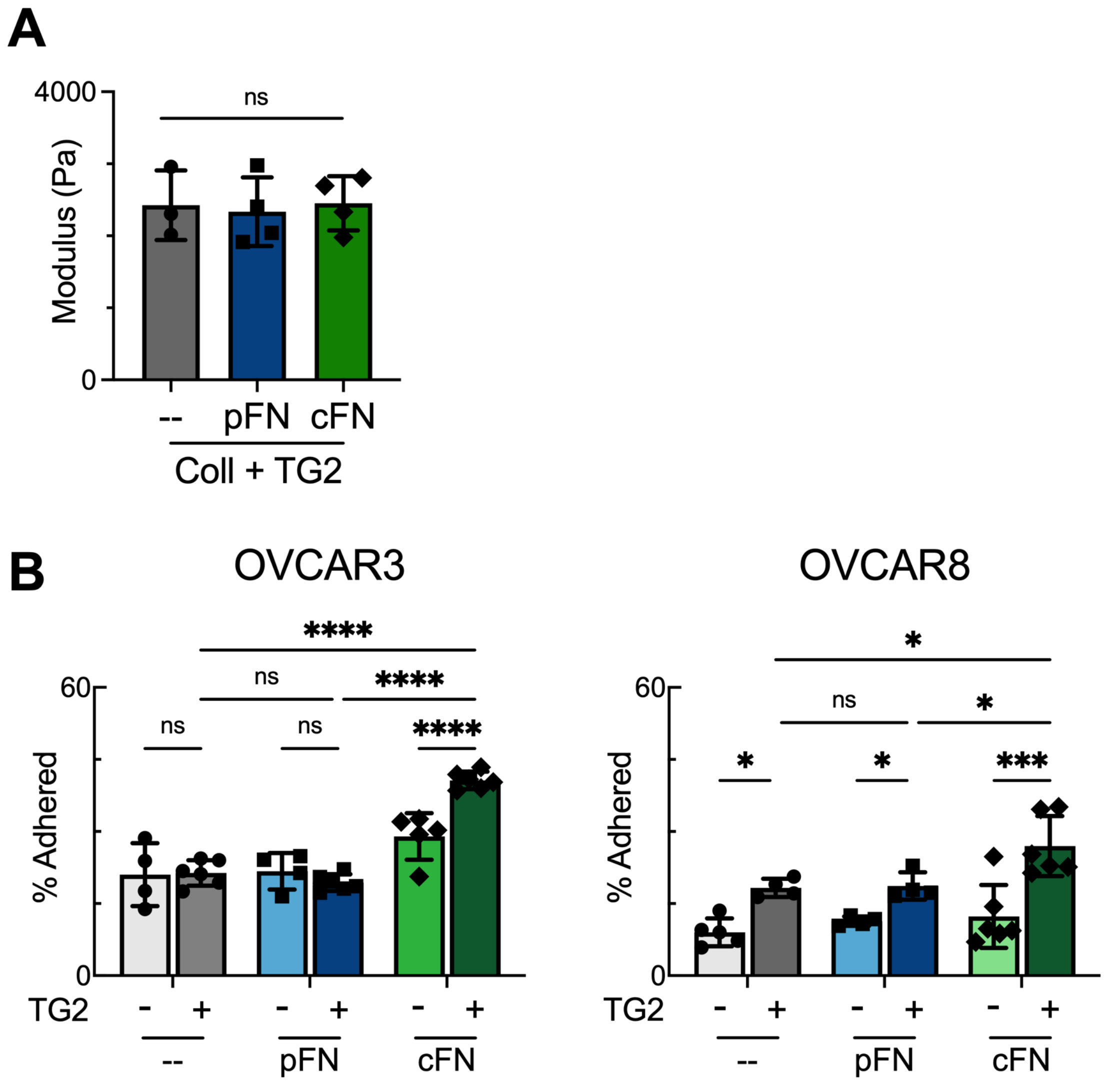


**Supplementary Figure 4.** Cross-linking gels with transglutaminase-2 increases cell adhesion. **A,** Elastic modulus is similar in transglutaminase cross-linked coll gels with various fibronectin additions. Each dot represents an individual gel. ns indicates p>0.05 by ANOVA. **B,** Quantification of OVCAR3 and OVCAR8 cells attached on various gel formulations. Each dot represents an individual gel, ns indicates p>0.05, * indicates p<0.05, *** indicates p<0.001, **** indicates p<0.0001 by mixed-effects model and Tukey. Data for gels without TG2 are the same as in Supplemental Figure 2C and the statistical comparison of -TG2 conditions are not included here.

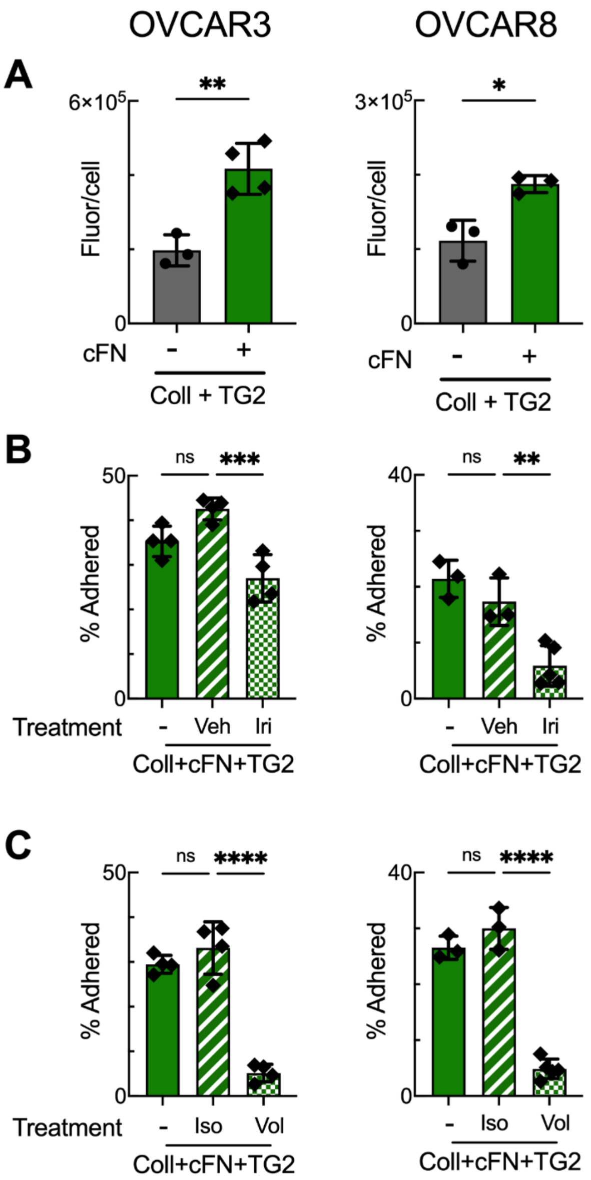


**Supplementary Figure 5.** Integrin engagement varies between different gel formulations. **A,** Quantification of the total fluorescence/cell of activated β1 integrin for OVCAR3 and OVCAR8 on coll+TG2 or coll+cFN+TG2 gels. Each dot represents the average across at least 3 fields of view for an individual gel, * indicates $p<0.05$, ** indicates $p<0.01$ by unpaired t-test. **B,C,** Quantification of the adhesion of OVCAR3 and OVCAR8 cells on coll+cFN+TG2 gels upon different treatments. -, no treatment control; Veh, DMSO control; Iri, irigenin; Iso, isotype control; Vol, volociximab. Each dot represents an individual gel, ns indicates $p>0.05$, ** indicates $p<0.01$, *** indicates $p<0.001$, **** indicates $p<0.0001$ by Tukey.

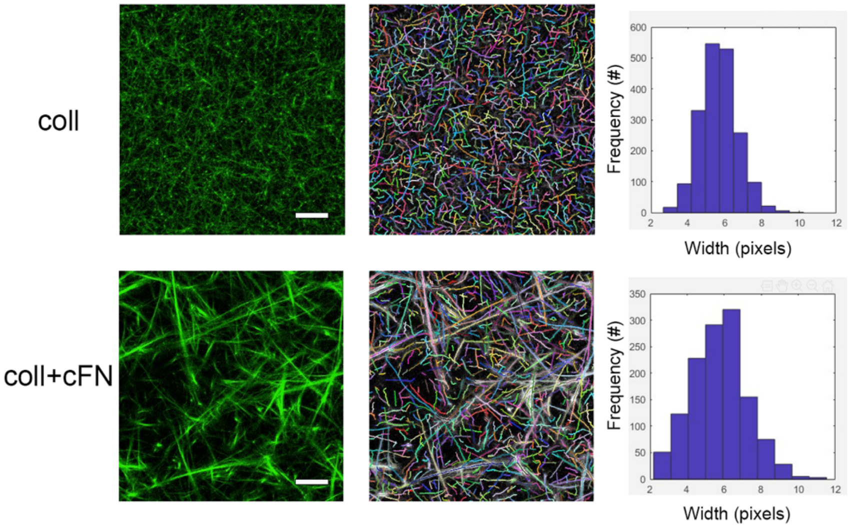


**Supplemental Figure 6.** Analysis of fiber metrics in coll and coll+cFN gels demonstrates a similar distribution of fiber widths. Shown are representative CNA35-EGFP stained coll and coll+cFN gels (left), CT-FIRE identified fibers (center), and distribution of fiber widths (right). Scale bar = 10 μm.

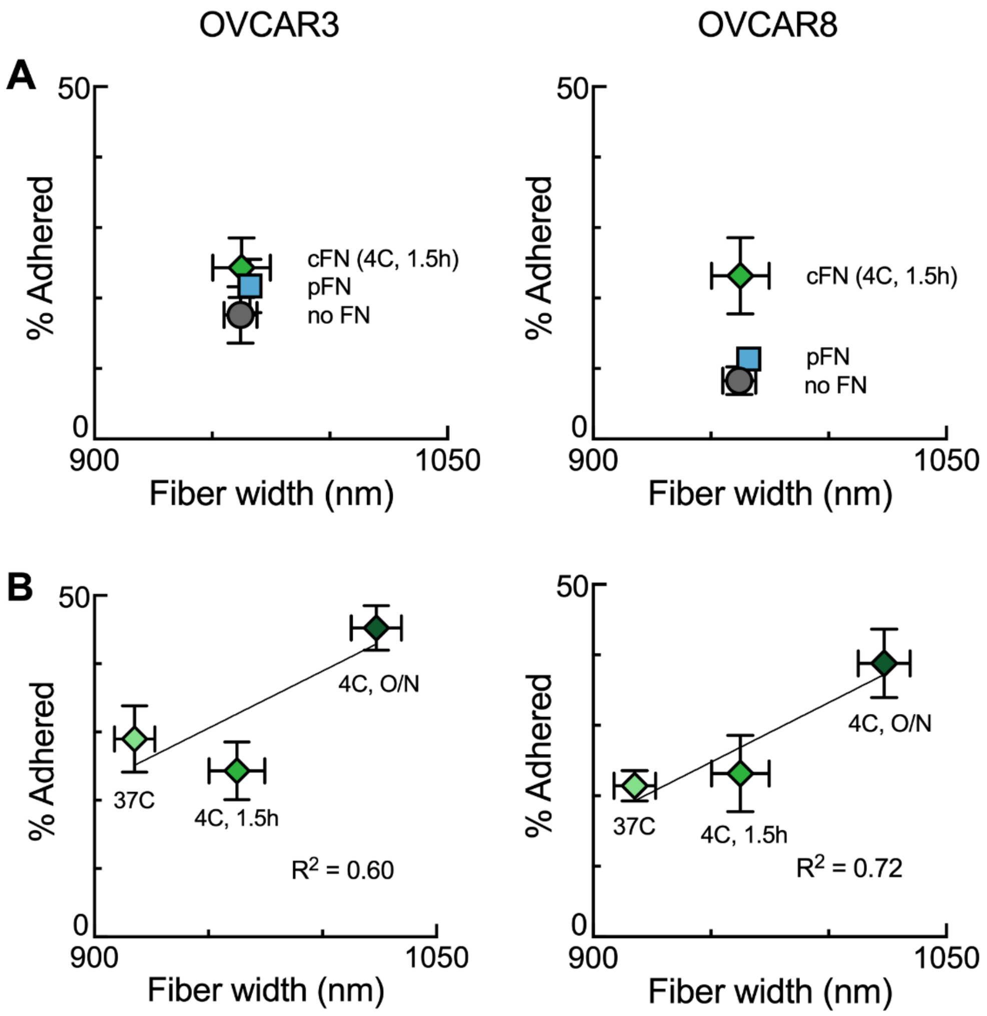


**Supplemental Figure 7.** HGSOC cell adhesion is impacted by fibronectin and fiber width. **A**, Quantification of mean fiber width and OVCAR3 or OVCAR8 cell adhesion for the gel conditions with varying FN additions are represented as average $\pm$ standard deviation, n = 3-5 gels per measurement. **B,** Quantification of fiber width and OVCAR3 or OVCAR8 cell adhesion for coll+cFN gels that were gelled with different conditions to result in a range of fiber widths are represented as average $\pm$ standard deviation, n = 3-6 gels per measurement.
